# Benchmarking challenging small variants with linked and long reads

**DOI:** 10.1101/2020.07.24.212712

**Authors:** Justin Wagner, Nathan D Olson, Lindsay Harris, Jennifer McDaniel, Ziad Khan, Jesse Farek, Medhat Mahmoud, Ana Stankovic, Vladimir Kovacevic, Byunggil Yoo, Neil Miller, Jeffrey A. Rosenfeld, Bohan Ni, Samantha Zarate, Melanie Kirsche, Sergey Aganezov, Michael Schatz, Giuseppe Narzisi, Marta Byrska-Bishop, Wayne Clarke, Uday S. Evani, Charles Markello, Kishwar Shafin, Xin Zhou, Arend Sidow, Vikas Bansal, Peter Ebert, Tobias Marschall, Peter Lansdorp, Vincent Hanlon, Carl-Adam Mattsson, Alvaro Martinez Barrio, Ian T Fiddes, Chunlin Xiao, Arkarachai Fungtammasan, Chen-Shan Chin, Aaron M Wenger, William J Rowell, Fritz J Sedlazeck, Andrew Carroll, Marc Salit, Justin M Zook

## Abstract

Genome in a Bottle (GIAB) benchmarks have been widely used to help validate clinical sequencing pipelines and develop new variant calling and sequencing methods. Here, we use accurate linked reads and long reads to expand the prior benchmarks in 7 samples to include difficult-to-map regions and segmental duplications that are not readily accessible to short reads. Our new benchmark adds more than 300,000 SNVs, 50,000 indels, and 16 % new exonic variants, many in challenging, clinically relevant genes not previously covered (e.g., *PMS2*). For HG002, we include 92% of the autosomal GRCh38 assembly, while excluding problematic regions for benchmarking small variants (e.g., copy number variants and reference errors) that should not have been in the previous version, which included 85% of GRCh38. By including difficult-to-map regions, this benchmark identifies eight times more false negatives in a short read variant call set relative to our previous benchmark.We have demonstrated the utility of this benchmark to reliably identify false positives and false negatives across technologies in more challenging regions, which enables continued technology and bioinformatics development.

## Introduction

Advances in genome sequencing technologies have continually transformed biological research and clinical diagnostics, and benchmarks have been critical to ensure the quality of the sequencing results. The Genome in a Bottle Consortium (GIAB) developed extensive data^1^ and widely used benchmark sets to assess accuracy of variant calls resulting from human genome sequencing.^2–4^ To use these benchmarks, the Global Alliance for Genomics and Health (GA4GH) Benchmarking Team develop tools and best practices for benchmarking.^5^ These benchmarks and benchmarking tools helped enable the development and optimization of new technologies and bioinformatics approaches, including linked reads,^6^ highly accurate long reads,^7^ deep learning-based variant callers,^8,9^ graph-based variant callers,^10^ and *de novo* assembly.^11,12^ As these new technologies and methods accessed increasingly challenging regions of the genome, studies highlighted many known medically relevant genes that were excluded from these previous benchmarks.^7,13–15^ These studies demonstrated the need for improved benchmarks covering segmental duplications, the Major Histocompatibility Complex (MHC), and other challenging regions. A separate synthetic diploid benchmark was generated from assemblies of error-prone long reads for two haploid hydatidiform mole cell lines, but this had limitations both in terms of cell line availability and small indel errors due to the high error rate of the long reads.^16^

Many of the difficult regions of the genome lie in segmental duplications and other repetitive elements. Linked reads were shown to have the potential to expand the GIAB benchmark by 68.9 Mbp to some of these segmental duplications.^6^ A circular consensus sequencing (CCS) method was recently developed that enables highly accurate 10 kb to 20 kb long reads.^7^ This technology identified a few thousand likely errors in the GIAB benchmark, mostly in LINEs. It also had >400,000 variants in regions mappable with long reads but outside the benchmark, and it covered many difficult-to-map, medically-relevant genes that are challenging to call using either short reads or lower accuracy long reads. GIAB recently used these data to produce a local diploid assembly-based benchmark for the highly polymorphic MHC region.^17^

Here, we use linked reads and long reads to expand GIAB’s benchmark to include challenging genomic regions for the GIAB pilot genome NA12878 and the GIAB Ashkenazi and Han Chinese trios from the Personal Genome Project, which are more broadly consented for genome sequencing and commercial redistribution of reference samples.^18^ We more carefully exclude segmental duplications that are copy number variable in the GIAB samples ^19^ or missing copies in GRCh37 or GRCh38,^20,21^ because these currently cannot be reliably benchmarked for small variants. We also refined the methods used to produce the diploid assembly-based MHC benchmark^17^ to include most of the MHC region in each member of the trio. We show that the new benchmark reliably identifies false positives and false negatives across a variety of short-, linked-, and long-read technologies. The benchmark has already been used to develop and demonstrate new variant callers in the precisionFDA Truth Challenge V2.^22^

## Results

### New benchmark covers more of the reference, including many segmental duplications

GIAB previously developed an integration approach to combine results from different sequencing technologies and analysis methods, using expert-driven heuristics and features of the mapped sequencing reads to determine at which genomic positions each method should be trusted. This integration approach excludes regions where all methods may have systematic errors or locations where methods produce different variants or genotypes and have no evidence of bias or error. While the previous version (v3.3.2) primarily used a variety of shor-tread sequencing technologies and excluded most segmental duplications,^4^ our new HG002 v4.2.1 benchmark adds long- and linked-reads to cover 6% more of the autosomal assembled bases for both GRCh37 and GRCh38 than v3.3.2 **(Table 1)**. Median coverage by linked- and long-read datasets for each genome is in **Supplementary Table 1.** We also replace the mapping-based benchmark with assembly-based benchmark variants and regions in the MHC.^17^ v4.2.1 includes more than 300,000 new SNVs and 50,000 INDELs compared to v3.3.2. In **Methods,** we detail the creation of the v4.2.1 benchmark, including using the new long- and linked-read sequencing data in the GIAB small variant integration pipeline, and identifying regions that are difficult to benchmark, including potential large duplications in HG002 relative to the reference as well as problematic regions of GRCh37 or GRCh38.

**Table 1:** Summary comparison between v3.3.2 and v4.2.1 HG002 benchmark set for chromosomes 1 to 22 in GRCh37 and GRCh38, including inclusion of segmental duplications (Seg Dups) and regions that appear similar to short reads (i.e., “low mappability” regions where 100 bp read pairs have <=2 mismatches and <=1 indel difference from another region of the genome).

| Reference Build | Benchmark Set | Reference Included | SNVs | Indels | Base pairs in Seg Dups and low mappability |
| --- | --- | --- | --- | --- | --- |
| GRCh37 | v3.3.2 | 87.8% | 3,048,869 | 464,463 | 57,277,670 |
| GRCh37 | v4.2.1 | 94.1% | 3,353,881 | 522,388 | 133,848,288 |
| GRCh38 | v3.3.2 | 85.4% | 3,030,495 | 475,332 | 65,714,199 |
| GRCh38 | v4.2.1 | 92.2% | 3,367,208 | 525,545 | 145,585,710 |

Many of the benchmark regions new to v4.2.1 are in segmental duplications and other regions with low mappability for short reads **(Figure 1, Supplementary Figure 1,** and **Table 1).** GRCh38 has 270,860,615 bases in segmental duplications and low mappability regions (regions difficult to map with paired 100 bp reads) on chromosomes 1 to 22, including modeled centromeres. v4.2.1 covers 145,585,710 (53.7%) of those bases while v3.3.2 covers 65,714,199 (24.3%) bases. However, v4.2.1 still excludes some difficult regions and structural variants; of the bases in GRCh38 chromosomes 1-22 not covered by v4.2.1, segmental duplication and low mappability regions account for 56.4% of those bases.

**Figure 1:**
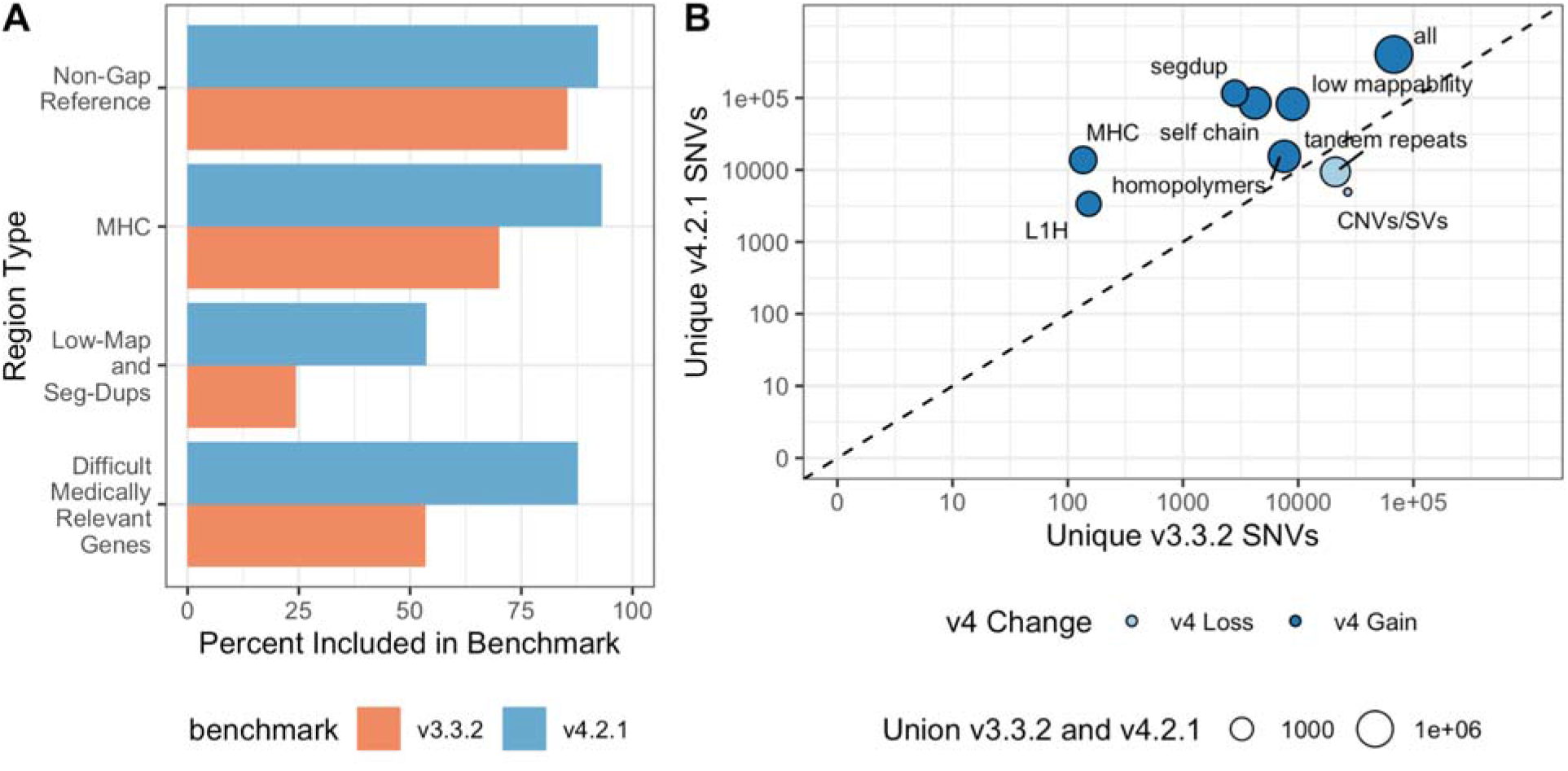
New benchmark set includes more of the reference genome and more SNVs and indels. (A) Percent of the genomic region that is included by v3.3.2 and v4.2.1 of all non-gap, autosomal GRCh38 bases; MHC; low mappability regions and segmental duplications; and 159 difficult-to-map, medically-relevant genes described previously. (B) The number of unique SNVs by genomic context. Circle size indicates the total number of SNVs in the union of v3.3.2 and v4.2.1. Circles above the diagonal indicate there is a net gain of SNVs in the newer benchmark, and circles below the diagonal indicate a net loss of SNVs in the newer benchmark.

To identify the types of genomic regions where v4.2.1 gains and loses benchmark variants relative to v3.3.2, we compared the variant calls in v4.2.1 vs. v3.3.2 and used the v2.0 GA4GH/GIAB stratification files.^22^ **Figure 1B** highlights stratified genomic regions with the largest SNV gains and losses in v4.2.1 vs. v3.3.2 (the full table is available in **Supplementary Table 2).** As expected, the inclusion of linked- and long-reads leads to more variants in v4.2.1 than v3.3.2 in segmental duplications, self chains, the MHC region, as well as other regions that are difficult to map with short reads. The gain in v4.2.1 relative to v3.3.2 is lower in tandem repeats and homopolymers because v4.2.1 excludes any tandem repeats and homopolymers not completely included by the benchmark regions. Partially included tandem repeats and homopolymers in v3.3.2 caused some errors in benchmarking results when v3.3.2 missed variants in the repeat but outside the benchmark regions, so partially included repeats were completely excluded in v4.2.1.

In addition to including more difficult regions, v4.2.1 also corrects or excludes errors in v3.3.2. In previous work, variants called from PacBio HiFi were benchmarked against v3.3.2, and 60 SNV and indel putative false positives were manually curated, which identified 20 likely errors in v3.3.2.^7^ All 20 errors were corrected in the v4.2.1 benchmark or removed from the v4.2.1 benchmark regions. Twelve of these errors in v3.3.2 result from short reads that were only from one haplotype, because reads from the other haplotype were not mapped due to a cluster of variants in a LINE; two of these v3.3.2 errors are excluded in v4.2.1, and 10 variants are correctly called in v4.2.1 **(Supplementary Table 3).** In order to verify the new v4.2.1 variants incorrectly called by v3.3.2 in LINEs, we confirmed all 274 tested variants in 4 LINEs across the 7 samples using Long-range PCR followed by Sanger sequencing, as described in **Methods** and **Supplementary Table 5.**

### New benchmark includes additional challenging genes

To focus analysis on potential genes of interest, we analyzed inclusion of genes previously identified to have at least one exon that is difficult to map with short reads, which we call “difficult-to-map, medically-relevant genes”.^13^ v4.2.1 covers 88 % of the 10,009,480 bp in difficult-to-map, medically-relevant genes on primary assembly chromosomes 1-22 in GRCh38, much larger than the 54% covered by v3.3.2 **(Table 2,** with GRCh37 in **Supplementary Table 4).** 3,913,104 bp of the difficult-to-map, medically-relevant genes lie in segmental duplication or low mappability regions. The v4.2.1 benchmark includes 2,928,012 bp (74.8%) of those segmental duplications and low mappability regions while the v3.3.2 benchmark includes 208,882 bp (5.3%). Future work will be needed to include 49 of the 159 genes on chromosome 1-22 that still have less than 90% of the gene body included (Figure 2A and **Supplementary Figure 2),** such as a new assembly-based approach.^21^ For example, 5 genes that have potential duplications in HG002 were previously partially included in v3.3.2 but are excluded in v4.2.1 because new methods will be needed to resolve and represent benchmark variants in duplicated regions **(Figure 2B).** The medically-relevant gene *KIR2DL1* was partially included in v3.3.2 but is now completely excluded because the copy number variable KIR region is removed from the v4.2.1 benchmark regions. v4.2.1 also does a better job excluding regions that are duplicated in the benchmark sample relative to the reference, specifically because it excludes regions with higher than normal PacBio HiFi and/or ONT coverage **(Figure 3).** We detail the inclusion of each difficult-to-map, medically-relevant gene in **Supplementary Table 6.**

**Table 2:** Benchmark inclusion of 159 medically relevant genes on chromosomes 1-22 previously identified as difficult for short reads. bp included is the total number of bp included by each benchmark set and percent of bases included from the gene set.

| Benchmark Set | bp included | SNVs | INDELS |
| --- | --- | --- | --- |
| v3.3.2 | 5,362,837 (54%) | 6,242 | 943 |
| v4.2.1 | 8,786,005 (88%) | 10,175 | 1,469 |

**Figure 2:**
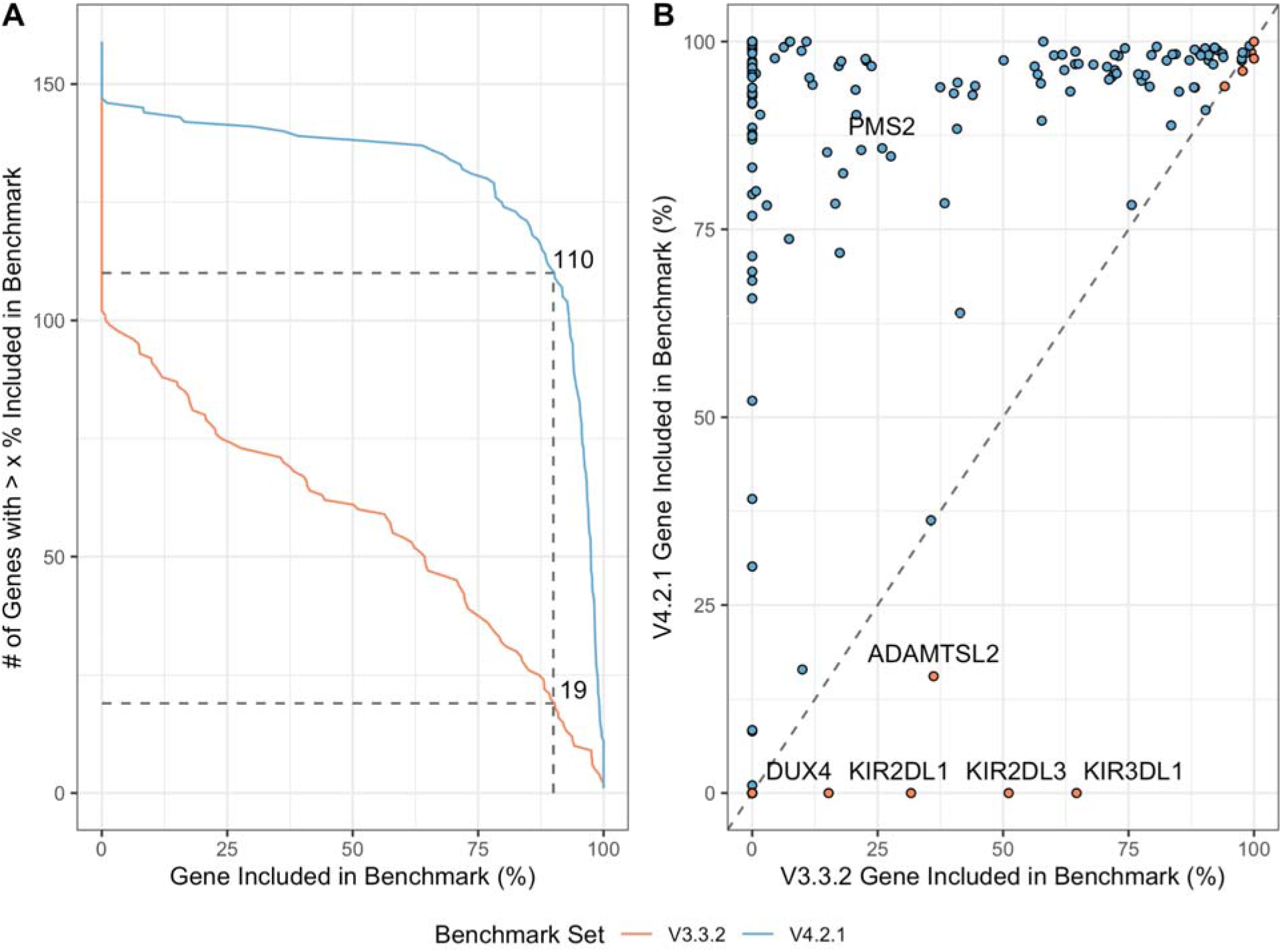
v4.2.1 includes many more difficult-to-map, medically-relevant genes. (A) Cumulative distribution for percent of each gene included in HG002 v4.2.1 benchmark regions for 159 autosomal difficult-to-map, medically-relevant genes. Dashed lines indicate that the number of genes included > 90% increased from 19 in v3.3.2 to 110 in v4.2.1. (B) Pairwise comparison of difficult-to-map, medically-relevant gene inclusion in benchmark set. Genes falling on the dashed line are similarly included by both benchmark sets, whereas genes above (red fill) or below (blue fill) the dashed line are included more by the v4.2.1 or v3.3.2 benchmark sets, respectively. The genes included more by v4.2.1 tend to be in segmental duplications and the smaller number of genes included more by v3.3.2 are mostly genes duplicated in HG002 relative to GRCh38 and should be excluded.

**Figure 3:**
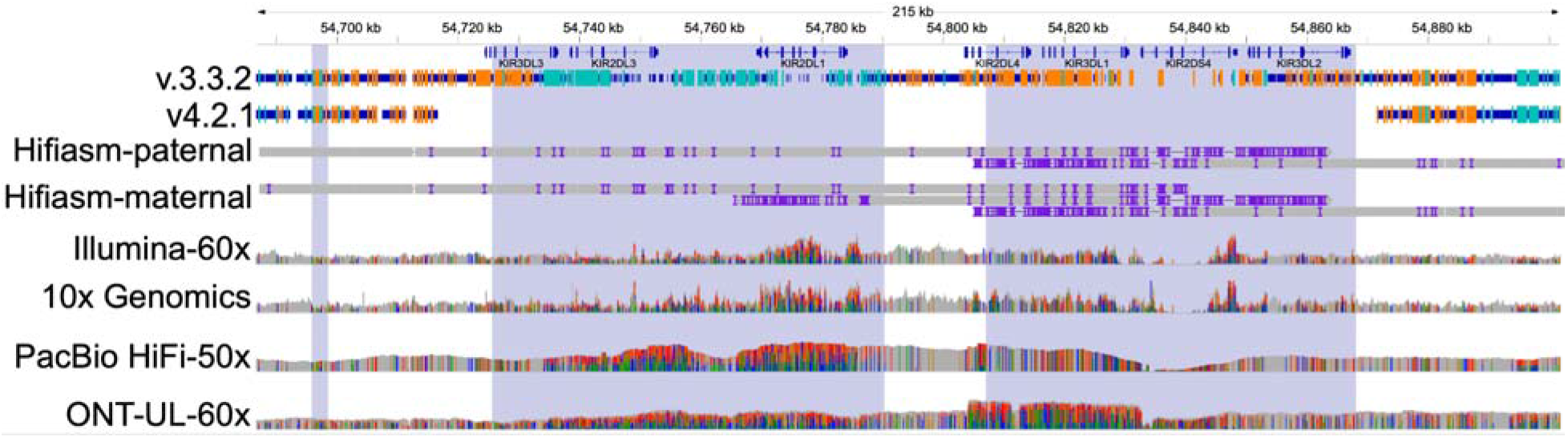
Genes in KIR locus are excluded in v4.2.1 due to duplication in HG002. edically-relevant genes in the KIR locus such as *KIR2DL1* were partially included in v3.3.2 with many erroneous variants, but are correctly excluded by v4.2.1 due to a likely duplication and other structural variation. Thick blue bars indicate regions included by each benchmark and orange and light blue lines indicate positions of homozygous and heterozygous benchmark variants, respectively. A duplication of part of this region, which is common in the population, is supported by higher than normal coverage and high variant density across all technologies, as well as alignment of multiple contigs from the maternal trio-based HG002 hifiasm assembly (Hifiasm-maternal). The region is very challenging to characterize and assemble accurately due to high variability and copy number polymorphisms in the population, as well as segmental duplications (shaded regions).

*PMS2* is an example of a medically important gene involved with DNA mismatch repair that is included more by v4.2.1 (85.6%) than by v3.3.2 (25.9%) for HG002 **(Figure 4).** Variant calling in *PMS2* is complicated by the presence of the pseudogene *PMS2CL*, which contains identical sequences in many of the exons of *PMS2* and is within a segmental duplication.^23^ Using Long Range PCR followed by Sanger sequencing, we confirmed 1,516 v4.2.1 benchmark variants in *PMS2* and 20 other difficult-to-map, medically-relevant genes across the 7 samples, and only 4 in PKD1 and 1 in FCGR2B were discordant with Sanger. The 5 discordant variants appeared to be clearly supported by short and long reads, and the reason for the discordant Sanger result was unclear. Detailed Sanger sequencing results for each gene and sample are in **Supplementary Table 5.**

**Figure 4:**
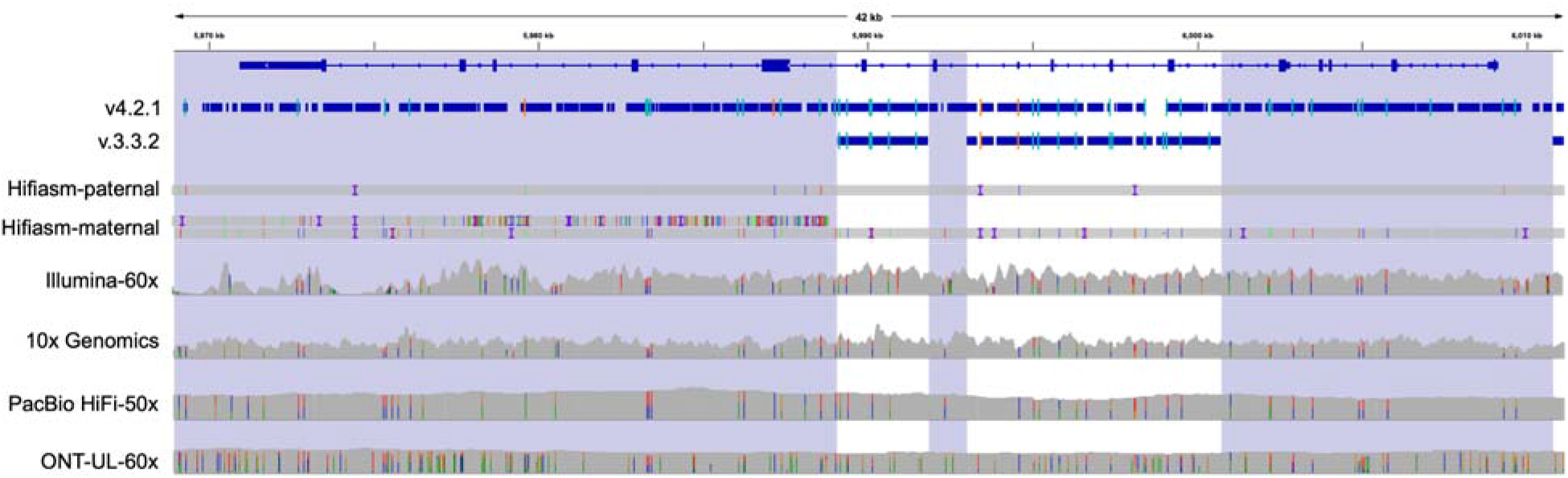
Difficult-to-map medically relevant gene *PMS2* is better included in v4.2.1. The medically-relevant gene *PMS2* is 85.6% included in the v4.2.1 benchmark regions while it is 25.9% included in v3.3.2 because segmental duplications (shaded regions) were largely excluded in previous benchmark versions. Thick blue bars indicate regions included by each benchmark and orange and light blue lines indicate positions of homozygous and heterozygous benchmark variants, respectively. This region is challenging for assembly-based approaches, and an extra contig from the maternal trio-based HG002 hifiasm assembly (Hifiasm-maternal) aligned to the left half of the gene due to mis-alignment or mis-assembly.

### Comparison to Platinum Genomes identifies fewer potential errors in v4.2.1

Platinum Genomes identified SNVs that were Mendelian inconsistent due to being called heterozygous in all 17 individuals in a pedigree with short read sequencing (“Category 1” errors).^24^ Some of these heterozygous calls result from regions duplicated in all individuals in the pedigree relative to GRCh37. Therefore, Category 1 SNVs matching SNVs in our benchmarks may identify questionable regions that should have been excluded from the benchmark regions. 326 Category 1 SNVs matched HG002 v4.2.1 SNVs, a decrease relative to the 719 Category 1 SNVs matching HG002 v3.3.2 SNVs. This suggests that v4.2.1 better excludes duplications in HG002 relative to the reference even as it expands into more challenging segmental duplication regions. However, the remaining 326 matching SNVs may be areas for future improvement in v4.2.1. Manual curation of 10 random SNVs in HG002 v4.2.1 that matched Category 1 variants showed 5 were in possible duplications that potentially should be excluded, and 5 were in segmental duplication regions that may have been short read mapping errors or more complex variation in segmental duplications **(Supplementary Table 7).** Particularly, clusters of v4.2.1 variants matching Category 1 variants appeared to be likely errors in v4.2.1. We also compared the v4.2.1 HG001 benchmark to the 2017 hybrid short-read benchmark from Platinum Genomes, which uses an orthogonal approach based on including variants with genotypes phased as expected in the 17-member pedigree. The concordance between v4.2.1 and Platinum Genomes in the intersection of both benchmark regions was 99.96% on GRCh37 and GRCh38. Curation identified most differences as likely short-read mapping biases in Platinum Genomes, as 454 of 654 GIAB-specific and 1857 of 2203 Platinum Genomes-specific variants on GRCh37 fell in low mappability regions and segmental duplications. In addition, relative to the short read-based Platinum Genomes benchmark regions, the v4.2.1 benchmark regions have substantially fewer small gaps that can cause problems when benchmarking,^4^ so that the NG50 size of benchmark regions in v4.2.1 is more than two times greater than Platinum Genomes **(Supplementary Figure 5).**

### High Mendelian Consistency in Trio

To further evaluate the accuracy of the benchmark, we evaluated the Mendelian consistency of our v4.2.1 benchmark sets for the son, father, and mother in two trios from GIAB of Ashkenazi ancestry (HG002, HG003, and HG004) and Han Chinese ancestry (HG005, HG006, and HG007). In the intersection of the benchmark regions for the Ashkenazi trio, this evaluation identified 2,502 variants that had a genotype pattern inconsistent with Mendelian inheritance out of the 4,968,730 variants in at least one member of the trio (0.05%), slightly below the rate for v3.3.2 (2,494 out of 4,383,371, or 0.06%) on GRCh38. The Mendelian inconsistency rates for the GIAB Han Chinese trio were lower than the Ashkenazi trio, 821/4601643 (0.02%) for v4.2.1 and 790/4138328 (0.02%) for v3.3.2. We separately analyzed Mendelian inconsistent variants that were potential cell line or germline *de novo* mutations (that is, the son was heterozygous and both parents were homozygous reference), and those that had any other Mendelian inconsistent pattern (which are unlikely to have a biological origin). Out of 2,502 violations in HG002, 1,177⍰SNVs and 284⍰INDELs were potential *de novo* mutations, 67 more SNVs and 71 more INDELs than in v3.3.2.^4^ HG005 had only 162 potential *de novo* SNVs and INDELs. Following the manual inspection of ten random de novo SNVs in HG002, 10/10 appeared to be true *de novo*. After manual inspection of ten random de novo indels, 10/10 appeared to be true *de novo* indels in homopolymers or tandem repeats. The violations that were not heterozygous in the son and homozygous reference in both parents fell in a few categories: (1) clusters of variants in segmental duplications where a variant was missed or incorrectly genotyped in one individual, (2) complex variants in homopolymers and tandem repeats that were incorrectly called or genotyped in one individual, and (3) some overlapping complex variants in the MHC that were correctly called in the trio but the different representations were not reconciled by our method (even though we used a method that is robust to most differences in representation).^4,25^ We exclude all Mendelian inconsistencies that are not heterozygous in the son and homozygous reference in both parents from the v4.2.1 benchmark regions of each member of the trio, because most are unlikely to have a biological origin. Conservative paternal | maternal phasing for HG002 on GRCh38 was performed for the MHC using local diploid assembly, and outside the MHC using phasing that was consistent between trio analysis and integrated Strand-seq and PacBio HiFi phasing (1,812,845/2,449,937 heterozygous benchmark variants).

### Regions excluded from the benchmark

A critical part of forming a reliable v4.2.1 benchmark was to identify regions that should be excluded from the benchmark. In **Table 3 and Supplementary Figure 6,** we detail each region type that is excluded, the size of the regions, and reasons for exclusion. We describe how each region is defined in **Methods,** and **Supplementary Note 2** describes refinements to these excluded regions between the initial draft release and the v4.2.1 benchmark. These excluded regions fall in several categories: (1) the modeled centromere and heterochromatin in GRCh38 because these are highly repetitive regions and generally differ in structure and copy number between any individual and the reference; (2) the VDJ, which encodes immune system components and undergoes somatic recombination in B cells; (3) in GRCh37, regions that are either expanded or collapsed relative to GRCh38; (4) segmental duplications with greater than 5 copies longer than 10 kb and identity greater than 99 %, where errors are likely in mapping and variant calling, e.g., due to structural or copy number variation resulting in calling paralogous sequence variants;^26,27^ (5) potential large duplications that are in HG002 relative to GRCh37 or GRCh38; (6) putative insertions, deletions, and inversions >49bp in size and flanking sequence; (7) tandem repeats larger than 10,000 bp where variants can be difficult to detect accurately given the length of PacBio HiFi reads. As an example of the importance of carefully excluding questionable regions, when comparing variants from ultralong reads to v3.3.2, 74 % of the putative FPs in HG002 on GRCh38 fell outside the v4.2.1 benchmark regions (see **Supplementary Table 8 and Supplementary Table 9).** Many of these were in centromere regions that have very few benchmark variants but were erroneously included in the v3.3.2 short read-based benchmark, e.g., in chr20. Our new benchmark correctly excludes these regions from the benchmark because they cannot be confidently mapped with short-, linked-, or long-reads used to form the benchmark.

**Table 3:**
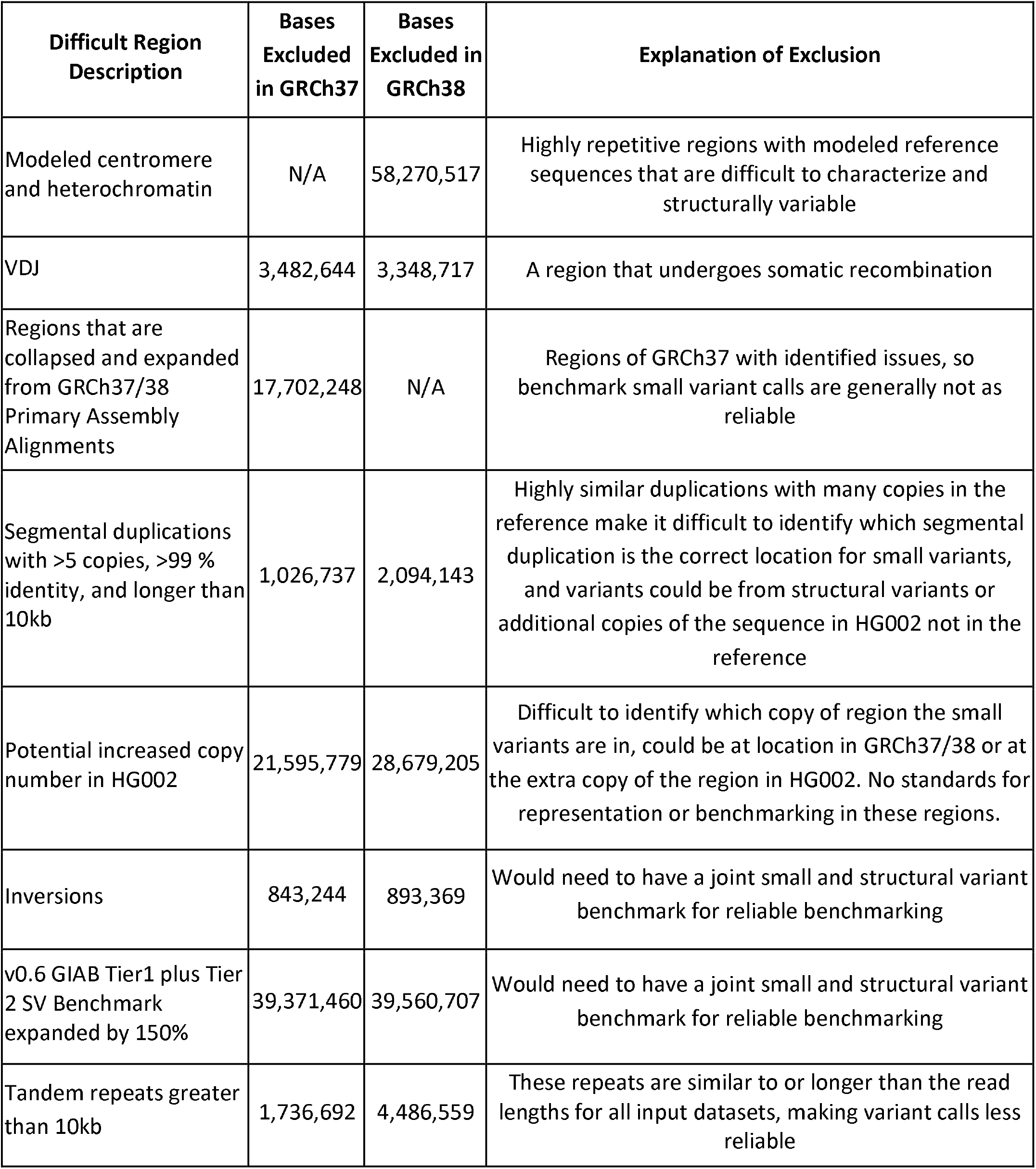
Base pairs overlapping different types of difficult regions that are excluded from all input callsets for HG002. The table shows progressive subtraction of other difficult regions so each row has all rows above it subtracted before calculating overlapping base pairs. In non-gap regions on chromosomes 1-22, there are 158,845,257 bp in GRCh37 and 202,943,679 bp in GRCh38 that are excluded by v4.2.1 (i.e., outside the v4.2.1 benchmark regions).

| **Difficult Region Description** | **Bases Excluded in GRCh37** | **Bases Excluded in GRCh38** | **Explanation of Exclusion** |
| --- | --- | --- | --- |
| Modeled centromere and heterochromatin | N/A | 58,270,517 | Highly repetitive regions with modeled reference sequences that are difficult to characterize and structurally variable |
| VDJ | 3,482,644 | 3,348,717 | A region that undergoes somatic recombination |
| Regions that are collapsed and expanded from GRCh37/38 Primary Assembly Alignments | 17,702,248 | N/A | Regions of GRCh37 with identified issues, so benchmark small variant calls are generally not as reliable |
| Segmental duplications with >5 copies, >99 % identity, and longer than 10kb | 1,026,737 | 2,094,143 | Highly similar duplications with many copies in the reference make it difficult to identify which segmental duplication is the correct location for small variants, and variants could be from structural variants or additional copies of the sequence in HG002 not in the reference |
| Potential increased copy number in HG002 | 21,595,779 | 28,679,205 | Difficult to identify which copy of region the small variants are in, could be at location in GRCh37/38 or at the extra copy of the region in HG002. No standards for representation or benchmarking in these regions. |
| Inversions | 843,244 | 893,369 | Would need to have a joint small and structural variant benchmark for reliable benchmarking |
| v0.6 GIAB Tier1 plus Tier 2 SV Benchmark expanded by 150% | 39,371,460 | 39,560,707 | Would need to have a joint small and structural variant benchmark for reliable benchmarking |
| Tandem repeats greater than 10kb | 1,736,692 | 4,486,559 | These repeats are similar to or longer than the read lengths for all input datasets, making variant calls less reliable |

### Evaluation and manual curation demonstrates reliability of benchmark

GIAB has established a community evaluation process for draft benchmarks before the official release.^3^ GIAB recruited volunteer experts in particular variant calling methods to follow the GA4GH Benchmarking Team’s Best Practices^5^ to compare a variety of query variant call sets to the draft benchmarks. We performed the community evaluation on v4.1 for HG002. Based on this evaluation, we made small improvements to generate v4.2.1 for HG002, as well as for the other 6 samples **(Supplementary Note 2).** v4.2.1 is the version described in the rest of this manuscript for all samples.

Query call sets for the final evaluation performed on v4.1 represented a broad range of sequencing technologies and bioinformatics methods **(Supplementary Table 10** and **Supplementary Note 1).** Each callset developer curated a random selection of FPs and FNs to ensure the benchmark reliably identifies errors in the query callset. Overall, we found that the benchmark was correct and the query callset was not correct in the majority of FP and FN SNVs and Indels **(Figure 5** with all curations in **Supplementary Table 11).** Overall, 433 of 452 (96%) curated FP and FN SNVs and INDELs inside v3.3.2 benchmark regions and 422 of 463 (91%) outside v3.3.2 benchmark regions were determined to be correct in the v4.1 benchmark. Some technologies/variant callers, particularly deep learning-based variant callers using HiFi data, had more sites where it was unclear if the benchmark was correct or the query callset was correct. These sites tended to be near regions with complex structural variation or places that appeared to be inside potential large duplications in HG002 but were not identified in our CNV approaches. In general, most sites that were not clearly correct in the benchmark and wrong in the query were in regions where the answer was unclear with current technologies **(Figure 5B).** For example, the v4.1 benchmark correctly excludes much of the questionable region in **Supplementary Figure 7,** but still includes some small regions where there may be a duplication and the variant calls both in the benchmark and the query are questionable. Future work will be aimed at developing a new benchmark in the small fraction of questionable regions, but these evaluations demonstrate the new benchmark reliably identifies both FPs and FNs across a large variety of variant callsets, including those based on short, linked, and long reads, as well as mapping-based, graph-based, and assembly-based variant callers.

**Figure 5:**
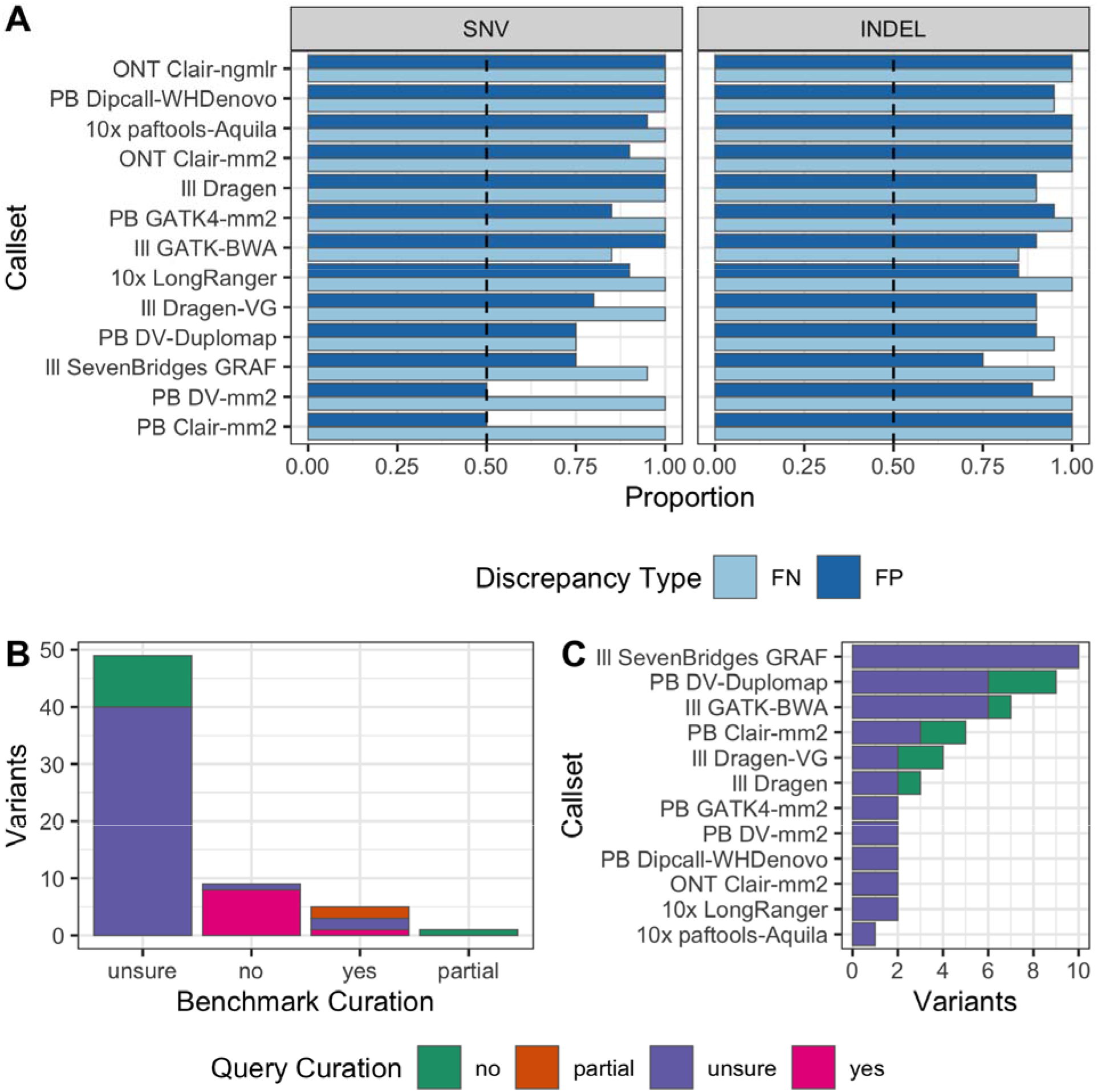
Summary of manual curations from the evaluation of the v4.1 benchmark, demonstrating it reliably identifies FPs and FNs in 10 callsets from different technologies and variant callers. (A) For each callset, we curated 20 FPs and 20 FNs, and this shows the proportion of curated FP and FN variants where the benchmark set was correct and the query callset was incorrect. The dashed black line indicates the desired majority threshold, 50%. Half of the curated variants were from GRCh37 and half were from GRCh38. (B) Breakdown of the total number of variants by manual curation category, excluding variants from panel A where the benchmark was deemed correct and query incorrect, showing most of these sites were difficult to curate. (C) Benchmark unsure variants by callset. Technology abbreviations are: ONT=Oxford Nanopore, PB=PacBio HiFi, Ill=Illumina PCR-free, 10x=10x Genomics

### New benchmark regions are enriched for false negatives

We demonstrate the benchmarking utility of v4.2.1 by comparing an example query call set to the new and old benchmark sets for HG002. For a standard short read-based call set (Ill GATK-BWA in **Figure 5),** the number of SNVs missed (even when including filtered variants) was 8.5 times higher when benchmarking against v4.2.1 than against v3.3.2 (16,615 vs. 1,960). The difference is largely due to false negative SNVs in regions of low mappability and segmental duplications with 15,220 in v4.2.1 vs. 1,381 in v3.3.2. When counting conservatively filtered SNVs as false negatives, v4.2.1 detected 71,165 more errors (183,568 in v4.2.1 vs. 112,403 in v3.3.2), similar to the increases seen with the noisy long-read-based syndip benchmark relative to v3.3.2.^16^ Also similar to syndip, the number of false positive SNVs was higher for v4.2.1 (25,328) than v3.3.2 (13,788) before conservative filtering. However, the number of false positive SNVs was actually lower for v4.2.1 (1,539) than v3.3.2 (2,370) after conservative filtering, likely due to removal of potential structural and copy number variants in v4.2.1. Relative to syndip, v4.2.1 for HG002 covers about 1 % fewer autosomal bases in GRCh38 but 16 % more bases in regions of low mappability and segmental duplications. Comparison of the results from the first and second precisionFDA challenges (based on v3.2 and v4.2, respectively), demonstrated similar changes in performance when expanding the benchmark; the combined false positive and false negative rates increased as much as 10-fold when the winners of the first challenge were benchmarked against v4.2.^22^ The more challenging variants and regions included in v4.2.1 enable further optimization and development of variant callers in segmental duplications and low mappability regions.

## Discussion

We present the first diploid small variant benchmark that uses short-, linked-, and long-reads to confidently characterize a broad spectrum of genomic contexts, including non-repetitive regions as well as repetitive regions such as many segmental duplications, difficult to map regions, homopolymers, and tandem repeats. We demonstrated that the benchmark reliably identifies false positives and false negatives in more challenging regions across many short-, linked-, and long-read technologies and variant callers based on traditional methods, deep learning,^8,9^ graph-based references,^10^ and diploid assembly.^12^ The benchmark was used in the precisionFDA Truth Challenge V2 held in 2020. This challenge focused on difficult regions not covered well by the v3.2 benchmark used in the first Truth Challenge in 2016, and SNV error rates of the winners of the first Truth Challenge increase by as much as 10-fold when evaluated against the v4.2 benchmark compared to the v3.2 benchmark.^22^

We designed this benchmark to cover as much of the human genome as possible with current technologies, as long as the benchmark genome sequence is structurally similar to the GRCh37 or GRCh38 reference. As a linear reference-based benchmark, it has advantages over global *de novo* assembly-based approaches by using reference information to resolve highly homozygous regions and some of the segmental duplications and other repeats where our samples are similar to the reference assembly. This reference-based approach enables users to take advantage of the suite of benchmarking tools developed by the Global Alliance for Genomics and Health Benchmarking Team, including sophisticated comparison of complex variants, standardized performance metrics, and stratification by variant type and genome context.^5^ However, our approach necessitates carefully excluding regions where our reference samples differ structurally from GRCh37 or GRCh38 due to errors in the reference, gaps in the reference, CNVs, or SVs. Developing benchmarks in these regions will require the development of methods to characterize these regions with confidence (e.g., using diploid assembly), standards for representing variants in these regions, and benchmarking methodology and tools. For example, for variants inside segmental duplications for which the individual has more copies than the reference, methods are actively being developed to assemble these regions,^26,28^ but no standards exist for representing which copy the variants fall on or how to compare to a benchmark.

It is critical to understand the limitations of any benchmark. Because our current benchmark excludes regions that structurally differ from GRCh37 or GRCh38, it will not identify deficiencies in mapping-based approaches that rely on these references nor highlight advantages of assembly-based approaches that do not rely on these references. While we have tried to exclude all regions where our samples differ structurally from the reference, some regions with gains in copy number remain, particularly in segmental duplications where these are more challenging to identify. Similarly, we may not exclude all inversions, particularly those mediated by segmental duplications. In addition, the benchmark still excludes many indels between 15 bp and 50 bp in size. Although we have significantly increased our coverage of difficult-to-map, medically-relevant genes, more work remains. Many of these genes are excluded due to putative SVs or copy number gains, and future work will be needed to understand whether these are true SVs or copy number gains, and if so, how to fully characterize these regions.

We expect that future benchmarks will increasingly use highly contiguous diploid assembly to access the full range of genomic variation. Our current benchmark is helping enable this transition by identifying opportunities to improve assemblies in the genome regions that are structurally similar to GRCh37 and GRCh38.

## Acknowledgments

We thank the Genome in a Bottle Consortium for ongoing feedback and discussions about the benchmark. We thank participants in the precisionFDA Truth Challenge V2 for helpful feedback about the v4.2 benchmarks for the trio. We thank Valerie Schneider for advice regarding alignments of GRCh38 to GRCh37. Chunlin Xiao was supported by the Intramural Research Program of the National Library of Medicine, National Institutes of Health. P.E. and T.M. acknowledge computational infrastructure provided by theCenter for Information and Media Technology at the University of Düsseldorf and funding from the German Research Foundation (grants 391137747 and 395192176), as well as support by the BMBF-funded de.NBI Cloud within the German Network for Bioinformatics Infrastructure (de.NBI) (031A537B, 031A533A, 031A538A, 031A533B, 031A535A, 031A537C, 031A534A and 031A532B). Certain commercial equipment, instruments, or materials are identified to specify adequately experimental conditions or reported results. Such identification does not imply recommendation or endorsement by the National Institute of Standards and Technology, nor does it imply that the equipment, instruments, or materials identified are necessarily the best available for the purpose.

## Competing Interests

AMW and WJR are employees and shareholders of Pacific Biosciences. AMB and ITF were employees and shareholders of 10X Genomics. FJS has received sponsored travel from Oxford Nanopore and Pacific Biosciences, and received a 2018 sequencing grant from Pacific Biosciences. AS and VK are employees of Seven Bridges. AC is an employee of Google Inc. and is a former employee of DNAnexus. AF and C-SC are employees of DNAnexus. SMES is an employee of Roche.

## Author Contributions

Conceptualization: JW, NDO, MS, JMZ

Data curation: JW, NDO

Formal Analysis - benchmark: JW, NDO, JMZ

Formal Analysis - phasing: JW, PE, TM, PL, VH, C-AM, JMZ

Methodology: JW, AC, JMZ

Project administration: JW, JMZ

Resources: CX

Software: JW

Supervision: JMZ

Validation: JW, LH, ZK, JF, MM, AS, VK, AMW, WJR, CX, BY, NM, BN, SZ, MK, SA, MS, GN, MB-B, WC, USE, CM, KS, XZ, AS, VB, AMB, ITF, AF, C-SC, FJS, AC, JMZ

Visualization: JW, NDO

Writing – original draft: JW, JMZ

Writing – review & editing: NDO, JAR, FJS

## STAR Methods

### Incorporating 10x Genomics and PacBio HiFi reads into small variant integration pipeline

v4.2.1 uses the same variant call sets as v3.3.2 from Complete Genomics,^29^ Illumina PCR-free (novoalign, GATK, and freebayes), and Illumina mate-pair (bwa mem, GATK, and freebayes).^30–32^ v4.2.1 uses 10x Genomics linked-read data and the variant calls from the LongRanger pipeline^6^, which makes calls both with and without using information from partitioning reads into haplotypes. In v3.3.2, we used conservative, haplotype-separated GATK calls from 10x Genomics, where calls were only made separately on each haplotype and coverage from both haplotypes was required. Also, v4.2.1 uses PacBio HiFi data using Sequel II with read lengths of 15kb and 20kb merged into a dataset that has approximately 47x to 68x coverage **(Supplementary Table 1)**, with variants subsequently called by GATK4 and DeepVariant.^7,8^ The 10x and PacBio HiFi data provide access to genomic regions that were previously inaccessible to short reads including segmental duplications. As shown in **Table 1** the number of base pairs in the benchmark that covers segmental duplications has increased with the incorporation of long- and linked-read data. **Table 4** lists the input data sets for the small variant integration pipeline to produce v4.2.1 for HG002.

**Table 4:**
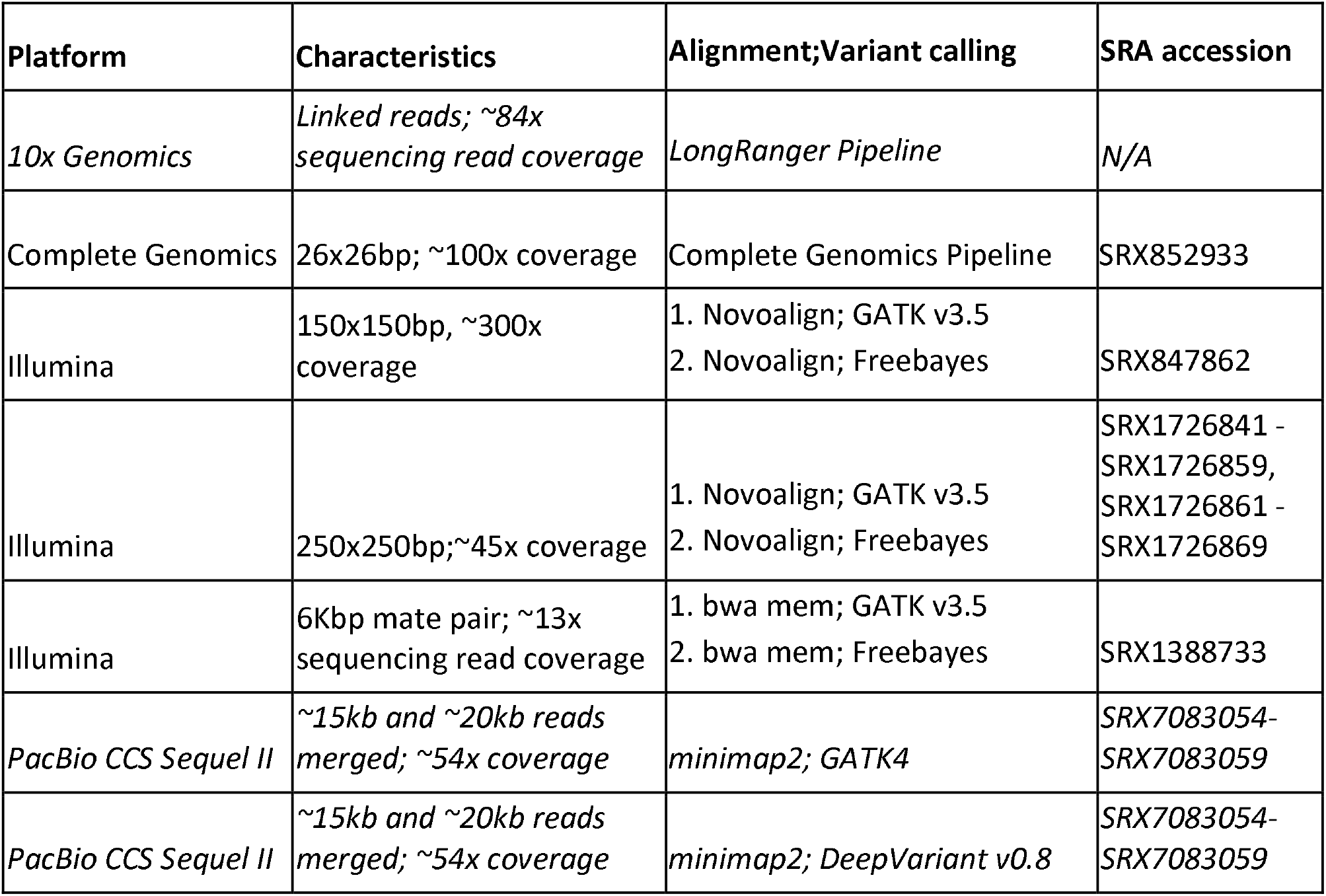
Integration Pipeline Data Set Attributes for HG002. Complete Genomics and Illumina data sets were the same as those used in v3.3.2 and were previously described.^1^ Italicized datasets from 10x Genomics and PacBio HiFi were new for v4.2.1.

| Platform | Characteristics | Alignment;Variant calling | SRA accession |
| --- | --- | --- | --- |
| *10x Genomics* | *Linked reads; ~84x sequencing read coverage* | *LongRanger Pipeline* | N/A |
| Complete Genomics | 26x26bp; ~100x coverage | Complete Genomics Pipeline | SRX852933 |
| Illumina | 150x150bp, ~300x coverage | 1. Novoalign; GATK v3.5<br>2. Novoalign; Freebayes | SRX847862 |
| Illumina | 250x250bp;~45x coverage | 1. Novoalign; GATK v3.5<br>2. Novoalign; Freebayes | SRX1726841 -<br>SRX1726859,<br>SRX1726861 -<br>SRX1726869 |
| Illumina | 6Kbp mate pair; ~13x sequencing read coverage | 1. bwa mem; GATK v3.5<br>2. bwa mem; Freebayes | SRX1388733 |
| *PacBio CCS Sequel II* | *~15kb and ~20kb reads merged; ~54x coverage* | *minimap2; GATK4* | *SRX7083054-*<br>*SRX7083059* |
| *PacBio CCS Sequel II* | *~15kb and ~20kb reads merged; ~54x coverage* | *minimap2; DeepVariant v0.8* | *SRX7083054-*<br>*SRX7083059* |

### Generating Callable Files with Haplotype-Separated BAMs

We use the CallableLoci utility in GATK3 to find regions with good coverage of high mapping quality reads. For PacBio HiFi and 10x Genomics read data, we use WhatsHap^33^ haplotag to partition reads by haplotype then use the bamtools filter function to generate separate BAM files for the two haplotypes. To partition reads by haplotype, we used a vcf that combined 10x linked read phasing with trio information described in the v0.6 structural variant benchmark paper.^3^ For CallableLoci with the unseparated BAM, we set the callable maxDepth threshold to 2 times the median coverage for VCF entries, then the minDepth threshold to 20. For the haplotype separated BAMs, we use median coverage for VCF as the maxDepth and 5 as the minDepth.

For PacBio HiFi, we first remove multi-allelic entries from the VCF and 50 bp on each side of the variant then remove RefCall entries with QUAL value below 40 along with 50 bp on each side of those variants. We then find callable regions for each haplotype BAM and the unseparated BAM then use bedtools multilntersectBed to find the union of those regions.

For 10x Genomics, we first remove filtered indels along with 50 bp on each side from its callable regions. Then we find callable regions on each haplotype and the unseparated BAM. After using multilntersectBed to find the union of those callable regions we subtract regions that were callable on one haplotype but not callable on the other haplotype.

### Python integration

We implemented the integration pipeline using Python as opposed to the Bash and Perl implementation for v3.3.2. The integration maintains a similar structure and we generated a DNAnexus applet to run on the same platform as v3.3.2. We updated the v4.2.1 pipeline to exclude all of a tandem repeat that is only partially covered by the benchmark regions. We also provide an option to not provide a callable file for given callsets, which for v4.2.1 we do not use callable regions for Ion Torrent or SOLiD. This results in a benchmark VCF that includes annotations for those technologies but variants are not excluded based on disagreement with their calls.

### Regions excluded from the benchmark

We determined regions to exclude from the benchmark where it was not currently possible to determine which technologies were correct due to the difficulty of resolving variation in that region. The difficult regions included those that had a structural variant identified in the GIAB SV v0.6 Benchmark, regions in which the HG002 sample had a copy variation compared to the reference, high depth and highly similar segmental duplications, tandem repeats > 10kb, regions that are collapsed and expanded from GRCh37/38 Primary Assembly Alignments, modeled centromere and heterochromatin, VDJ, and inversions. The bed files excluded from the benchmark are being made available in the v3.00 stratifications from GIAB under https://ftp-trace.ncbi.nlm.nih.gov/ReferenceSamples/giab/release/genome-stratifications/. We refined these regions with several rounds of internal and external evaluation on intermediate versions of the benchmark. We describe intermediate versions of the benchmark in **Supplementary Note 2.**

#### Modeled centromere and heterochromatin

We use the modeled centromere for GRCh38 from ftp://ftp-trace.ncbi.nlm.nih.gov/ReferenceSamples/giab/release/NA12878_HG001/NISTv3.3.2/GRCh38/supplementaryFiles/genomic_regions_definitions_modeledcentromere.bed and the heterochromatin region ftp://ftp-trace.ncbi.nlm.nih.gov/ReferenceSamples/giab/release/NA12878_HG001/NISTv3.3.2/GRCh38/supplementaryFiles/genomic_regions_definitions_heterochrom.bed.^34^

#### VDJ region subject to somatic recombination

We downloaded the UCSC genes tracks^35^ from http://hgdownload.cse.ucsc.edu/goldenPath/hg19/database/kgXref.txt.gz and selected entries with “abParts”. We then subset to chromosomes 2, 14, and 22 which contain the IGK, IGH, and IGL components that make up the VDJ recombination regions.

#### KIR region

v4.2.1 excludes the KIR region (chr19:54716689-54871732 in GRCh38 and 19:55228188-55383188 in GRCh37) because it is highly variable in copy number in the population, variant representation is challenging, and our current mapping-based methods have more errors in this region.

#### Regions that are collapsed and expanded from GRCh37/38 Primary Assembly Alignments

The GRC aligned GRCh37 to GRCh38 (excluding alts) with results available at: ftp://ftp.ncbi.nlm.nih.gov/pub/remap/Homo_sapiens/2.1/GCF_000001405.13_GRCh37/GCF_000001405.26_GRCh38/. We parsed the file GCF_000001405.13.xlxs for two Discrepancy values: (1) SP that denotes collapsed regions and (2) SP-only that denotes a region that was expanded between the reference versions.

#### Highly similar and high depth segmental duplications longer than 10kb

We begin with the segmental duplications track from UCSC^35^: http://hgdownload.cse.ucsc.edu/goldenPath/hg19/database/genomicSuperDups.txt.gz. We filter for entries larger than 10 kb and with identity > 99%. We then use bedtools genomecov to calculate segmental depth and subset to those greater than 5.

#### Potential copy number variation

We employed several approaches to find potential regions of large duplications in HG002 that are not in GRCh37 and GRCh38:

1. Short read and Long Read Intersection: We used mosdepth^36^ to find 1,000 bp windows that have higher than average coverage/2*2.5 in ONT and PacBio HiFi data. We intersected these regions with results from the CNV analysis tool, mrCaNaVar,^37^ on Illumina HiSeq 300x data (ftp://ftp-trace.ncbi.nlm.nih.gov/ReferenceSamples/giab/data/AshkenazimTrio/analysis/BilkentUni_IlluminaHiSeq_TARDIS_mrCaNaVar_05212019/AJtrio-HG002.hs37d5.300x.bam.bilkentuniv.052119.dups.bed.gz and ftp://ftp-trace.ncbi.nlm.nih.gov/ReferenceSamples/giab/data/AshkenazimTrio/analysis/BilkentUni_mrCaNaVaR_GRCh38_07242019/AJtrio-HG002.hg38.300x.bam.bilkentuniv.072319.dups.bed.gz).
2. Diploid Assemblies of HG002: We used SVRefine (https://github.com/nhansen/SVanalyzer) to align diploid assemblies to GRCh37/GRCh38 with bedgraph files that denote coverage of the reference by the number of contigs for the maternal and paternal haplotypes. We used bedtools unionBedGraphs and then found reference regions that are covered by 2 or more contigs in the union of haplotypes. We did this separately on a TrioCanu assembly using ONT,^38^ a Flye assembly using ONT,^39^ and a TrioCanu assembly of PacBio HiFi 15kb reads.^7^ We found an intersection across the three assemblies and subset to regions greater than 10 kb.
3. Elliptical Outlier Boundary with PacBio HiFi and ONT sequencing data: We used mosdepth to calculate coverage in 1,000 bp windows of the PacBio HiFi data and the ONT ultralong data set (ftp://ftp-trace.ncbi.nlm.nih.gov/ReferenceSamples/giab/data/AshkenazimTrio/HG002_NA24385_son/Ultralong_OxfordNanopore/guppy-V2.3.4_2019-06-26/ultra-long-ont_hs37d5_phased.bam and ftp://ftptrace.ncbi.nlm.nih.gov/ReferenceSamples/giab/data/AshkenazimTrio/HG002_NA24385_son/Ultralong_OxfordNanopore/guppy-V2.3.4_2019-06-26/ultra-long-ont_GRCh38_reheader.bam). We then found regions that had outlier coverage in PacBio HiFi and/or ONT. To do so, as described in the equations below, we (1) divided the PacBio HiFi coverage of each window by the median depth HiFi depth and squared it; (2) divided the ONT coverage of each window by the median depth ONT depth and squared it; (3) summed those values; and (4) took the square root of the sum. We found the third quartile and interquartile range for those transformed window coverage values. Finally, we found windows with coverage greater than the third quartile plus 1.5 times the IQR. In the equations below, WindowHiFiDepth, WindowONTDepth, and EllipticalValues are vectors, while MedianHiFiDepth, MedianONTDepth, and ThresholdEllipticalOutlier are scalars.

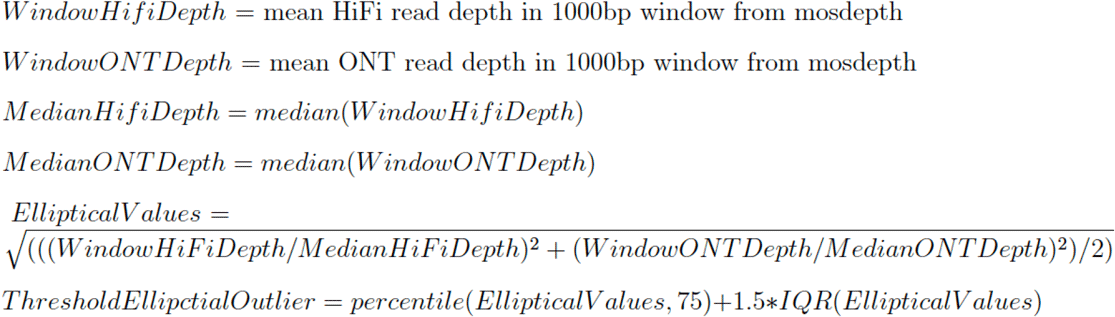

#### Inversions

We used SVrefine (github commit f0fb99969b6e239d1f49bc64a8f6cf5d52a2b88b) to call structural variants with, --maxsize 1000000 option. We then extracted inversions from the call set. Variants were merged with SVmerge (github commit aa8beb6f1cb5c539eea9f980ff30d2648caeee21), default maximum “distances”, which were 0.2 for all. SVrefine and SVmerge were from SVanalyzer (https://github.com/nhansen/SVanalyzer).

#### HG002 v0.6 GIAB Tier1 plus Tier 2 SV Benchmark expanded by 150%

We used bedtools^40^ slop with parameters -b -pct .25 to expand the GIAB v0.6 Structural Variant benchmark file: ftp://ftp-trace.ncbi.nlm.nih.gov/ReferenceSamples/giab/data/AshkenazimTrio/analysis/NIST_SVs_Integration_v0.6/HG002_SVs_Tier1plusTier2_v0.6.1.bed. This file defines regions in which calls with PASS in the FILTER field as well as regions should contain close to 100% of true insertions and deletions >=50 bp, with variants merged into a single region if they were within 1 kb.

#### SVs excluded from HG001 and HG003-HG007

Because we don’t have SV benchmarks for HG001 and HG003-HG007, we used pbsv (https://github.com/PacificBiosciences/pbsv) SVs >49 bp from PacBio HiFi data for HG001 and HG003-HG007, and well as svanalyzer and dipcall SVs >49 bp from trio-hifiasm assemblies of HG001 and HG005. We expanded these SVs to include overlapping tandem repeats and homopolymers and expanded the resulting regions by 25 % of the region size on each side with bedtools^40^ slop with parameters -b -pct .25.

#### Tandem Repeats greater than 10,000 bp

We took the union of SimpleRepeat dinucleotide, trinucleotide, and tetranucleotide STRs as well as RepeatMasker_LowComplexity, RepeatMasker_SimpleRepeats, and TRF_SimpleRepeats downloaded from UCSC Genome Browser. We then subset to tandem repeats longer than 10,000bp.

#### Reference assembly contigs shorter than 500,000 bp

We downloaded the gap track from UCSC^35^: ftp://hgdownload.cse.ucsc.edu/goldenPath/hg19/database/gap.txt.gz. Then subset to regions that are gap. We used bedtools complemented with GRCh37/GRCh38 to find contigs then subset to those less than 500 kb.

### Regions excluded for specific technologies

We exclude tandem repeats approximately larger than the read length from each method. Tandem repeats shorter than 51 bp were excluded from all technologies except Illumina PCR-free GATK, Complete Genomics, and PacBio HiFi DeepVariant. We excluded tandem repeats between 51 bp and 200 bp except for Illumina PCR-Free GATK and PacBio HiFi DeepVariant. Tandem repeats between 200 bp and 10,000 bp are excluded from all technologies except PacBio HiFi DeepVariant. Homopolymers greater than 6 bp were excluded from all technologies except Illumina PCR-free GATK, Complete Genomics, Ion Exome, PacBio HiFi DeepVariant. Imperfect homopolymers greater than 10 bp were excluded from all technologies except Illumina PCR-Free GATK. Low mappability regions that are difficult to map for short reads were excluded from all except 10x Genomics and PacBio HiFi. LINE:L1Hs greater than 500 bp were excluded except Illumina MatePair, 10x Genomics, and PacBio HiFi. Segmental duplications were excluded from all technologies except 10x Genomics and PacBio HiFi. Homopolymers were excluded from 10x Genomics and PacBio HiFi. Long homopolymers were included only for GATK based calls for PCR-Free data because GATK gVCF has low genotype quality score if reads do not totally encompass the homopolymer. Overall we trust homopolymers most from PCR-Free short reads. We visualize the regions excluded from each sequencing technology in **Figure 6.**

**Figure 6:**
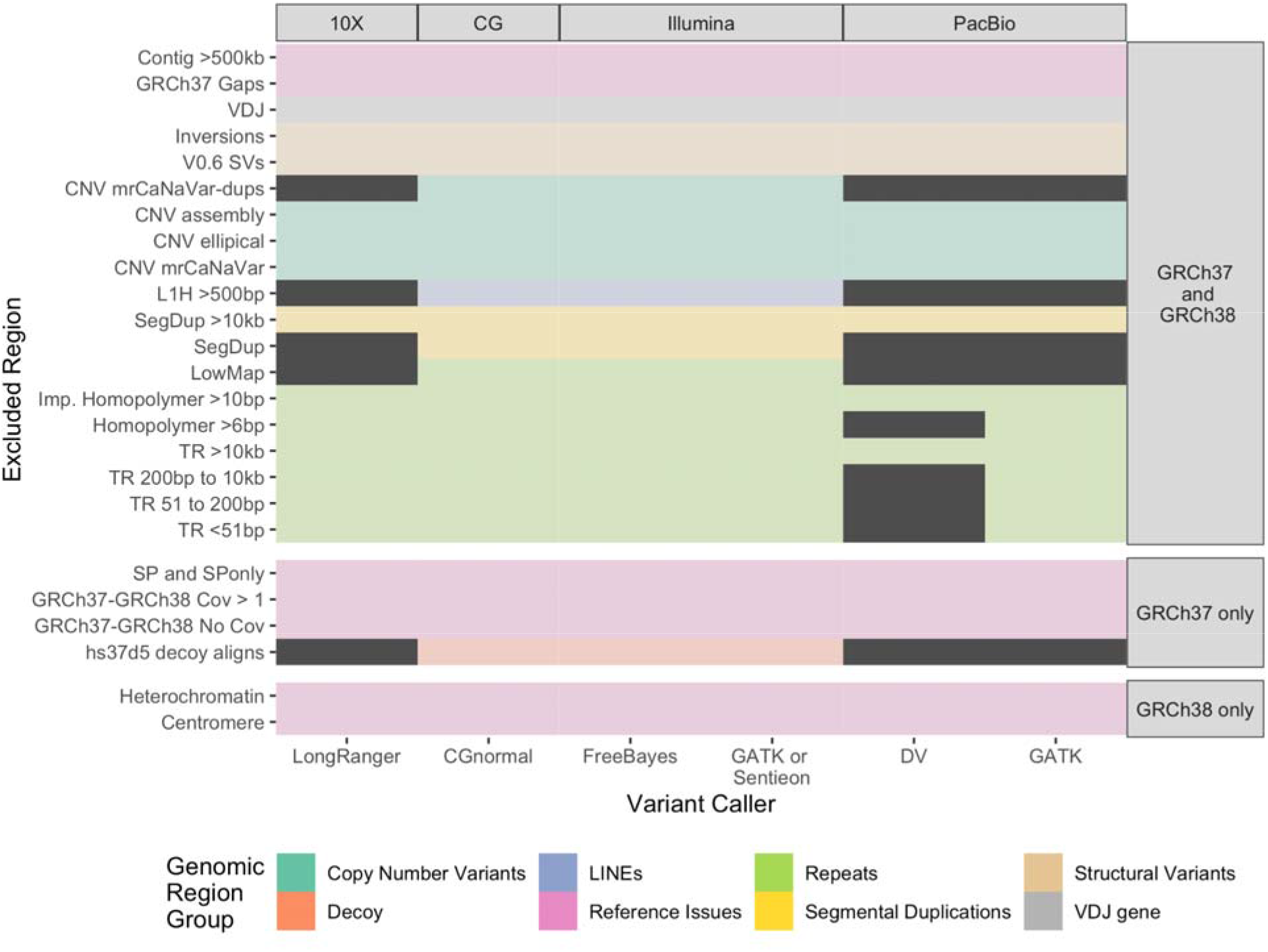
Genomic regions included by input variant callset. Genomic regions are excluded based on the biases of each technology that decrease reliability of variants in particular regions. Included regions are indicated by dark grey. Illumina PCR-Free includes both the high coverage HiSeq 300x and 2×250 HiSeq datasets. The PacBio HiFi dataset consists of 4 SMRT Cells of 15 kb libraries and 2 SMRT Cells of 20 kb libraries.

### Comparing v3.3.2 to v4.2.1

For HG002, we subset v3.3.2 variants to v3.3.2 benchmark bed and v4.2.1 variants to v4.2.1 benchmark bed and compared the benchmarks using hap.py with v2.0 of the GA4GH benchmarking stratifications (https://github.com/ga4gh/benchmarking-tools).^5^ To identify the types of genomic regions where v4.2.1 gains and loses benchmark variants relative to v3.3.2, we subset to stratifications with at least 1000 variants in v4.2.1, and sorted by the difference between the Precision and Recall metrics, which are measures of the fraction of extra variants in v3.3.2 and v4.2.1, respectively.

### Calculating difficult-to-map, medically-relevant genes coverage

We used the 193 clinically-relevant gene names that contained exons that are difficult to map with short reads from ^13^. We used Ensembl BioMart to retrieve Human Genes Build 99 with Gene Name, Start, End, and Chromosome (http://jan2020.archive.ensembl.org/biomart/martview/2c3a4b803e1a01b3b806829a466b3590).^41^ We used those results to find coordinates for the difficult-to-map, medically-relevant gene names, subset to genes on chromosomes 1-22, then used bedtools intersect with the v3.3.2 and v4.2.1 benchmark region files to find overlap.

### Evaluation of the benchmark

We used hap.py (https://github.com/Illumina/hap.py) following GA4GH best practices^5^ with HG002 v4.1 benchmark variants as the truth set, v4.1 benchmark bed as confident regions, and each of the 12 call sets as the query. We use the vcfeval engine for comparison.^25^

To evaluate the utility of the v4.1 benchmark, the GIAB community contributed 13 call sets from short-, linked-, and long-read technologies, and from mapping-, graph-, and assembly-based variant callers. We used hap.py to compare each input callset to v4.1 then asked collaborators to manually curate a small subset of the False Positive and False Negative sites with commands detailed in “Supplementary Materials - Benchmark Evaluations”. Collaborators evaluated 5 False Positive SNVs, 5 False Positive Indels, 5 False Negative SNVs, 5 False Negative Indels both inside and outside v3.3.2 along with 5 False Positive SNVs, 5 False Positive Indels, 5 False Negative SNVs, 5 False Negative Indels in the MHC for GRCh37. We generated IGV sessions with BAM files for Illumina HiSeq, 10x Genomics, PacBio HiFi 15kb & 20 kb merged, and ONT Ultralong^11^, then asked that the evaluators identify for each site if both alleles in the benchmark were correct and if both alleles in the query call set were correct.

### Long Range PCR Confirmation

We performed Long range PCR followed by Sanger sequencing for variants in LINEs and difficult-to-map, medically-relevant genes for all 7 samples. The difficult genes that were chosen for long-range PCR and Sanger sequence confirmation are potentially medically-relevant and have many characteristics that make them difficult to characterize, especially with short reads. We selected genes with previously published long range PCR assays. The first set of genes make up the RCCX complex, a segmental duplication that includes *TNXA*, *TNXB*, *C4A*, *C4B*, and *CYP21A2*.^42,43^ The similar sequences of these genes in close proximity makes them prone to rearrange, mutate and change the size of the complex as a whole, and they are linked to rare diseases that are inherited together at a higher rate than would be expected by chance.

Mutations in the *CYP2D6* gene can affect metabolism and bioactivation of many clinical drugs and the gene contains a polymorphic region.^44^ *DMBT1* has been identified as a candidate tumor suppressor for brain, gastrointestinal and lung cancers and contains highly repetitive sequence.^45^ Rare variants in the *HSPG2* gene are linked to cases of idiopathic scoliosis.^46^ *STRC* has a pseudogene with high genomic and coding sequence homology making it very difficult to characterize by normal short read sequencing methods.^47^ The *PMS2* gene has multiple pseudogenes, making it difficult to reliably detect mutations or characterize by sequencing.^23^ We additionally include v4.2.1 variants covered by the long range PCR assays designed for genes as described for the GIAB Challenging Medically Relevant Gene benchmark.^21^

Long range PCR was performed to amplify regions with variants in LINEs and difficult-to-map, medically-relevant genes. Primers for amplification of LINEs were designed with the Primer3Plus software.^48^ Other primers were sourced from literature. All long range primer sequences and references can be found in **Supplementary Table 12.** Long range PCR were performed with the PrimerSTAR GXL DNA Polymerase (Takara Bio, Mountain View, CA), and assays specific reaction components can be found in **Supplementary Table 13.** Long range PCR conditions varied by assay and can be found in **Supplementary Table 14.**

Sanger primers were designed using the Primer3Plus software.^48^ Primer sequences can be found in **Supplementary Table 12.** Long range PCR products were purified with ExoSAP-IT (Applied Biosystems, Foster City, CA). Sanger sequencing was performed with SimpleSeq Premixed Sequencing Kits (Eurofins Genomics, Louisville, KY) using 5 mL of the long range PCR amplicon and 5 mL of 3 mM primer. Sanger sequencing traces were aligned and analyzed with Geneious Prime (Biomatters, Inc., San Diego, CA).

### Phasing variant calls

To provide initial conservative phasing information for regions including the MHC and segmental duplications, the v4.2.1 benchmark vcf for HG002 on GRCh38 was phased in 3 ways. For the MHC, phasing was obtained from the fully phased local diploid assembly, using trio information to ensure it follows the paternal | maternal convention in the GT field. For the rest of the genome, we used phased heterozygous calls that were consistent in a single phase block for each chromosome between trio-based phasing and integrative phasing using Strand-seq and PacBio HiFi reads. The HG002 v4.2.1 benchmark variants were phased independently from the parental variants using integrative phasing.^49^ The integrative phasing approach combined local phase information from PacBio HiFi long-read alignments with global phase information obtained from Strand-seq short-read alignments to create whole-chromosome haplotypes for each individual. Method and implementation were applied as previously described^50^ with minor modifications: the GRCh38 assembly was used as reference for both PacBio HiFi long-read and Strand-seq short-read alignments, and the “--indels” option was added to the “whatshap phase” command line.

Additionally, for the children HG001, HG002, and HG005, we transferred paternal | maternal phasing from a dipcall^16^ vcf using a trio-hifiasm v0.11 assembly^51^ to v4.2.1 vcf of each individual. These draft phased vcfs are available under the SupplementaryFiles directory for HG001, HG002, and HG005 at https://ftp-trace.ncbi.nlm.nih.gov/ReferenceSamples/giab/release/.

### Data availability

Sequence data used is in Table 4, and is in SRA accessions SRX852933, SRX847862, SRX1726841 - SRX1726859, SRX1726861 - SRX1726869, SRX1388733, and SRX7083054 - SRX7083059. Aligned reads and other analyses from each sample are available at https://ftp-trace.ncbi.nlm.nih.gov/ReferenceSamples/giab/data/. The v4.2.1 benchmark vcf and bed files are available at: https://ftp-trace.ncbi.nlm.nih.gov/ReferenceSamples/giab/release/. GIAB’s v3.00 stratifications designed for use with the v4.2.1 benchmarks and the hap.py benchmarking tool, as well as the bed files excluded from the benchmark are being made available under https://ftp-trace.ncbi.nlm.nih.gov/ReferenceSamples/giab/release/genome-stratifications/.

### Code availability

Scripts for integrating candidate variants to form the benchmark set in this manuscript are available under https://github.com/jzook/genome-data-integration. Publicly available software used to generate input callsets and evaluation callsets is described in the methods and Supplementary Materials.

### Materials availability

DNA extracted from a single large batch of cells for 5 of the 7 genomes (HG001 to HG005) is publicly available in National Institute of Standards and Technology Reference Materials 8391 (HG001), 8392 (HG002-HG004), 8393 (HG005), and 8398 (HG001). DNA for HG001 to HG005, as well as HG006 and HG007, are extracted from publicly available cell lines GM12878 (RRID:CVCL_7526), GM24385 (RRID:CVCL_1C78), GM24149 (RRID:CVCL_1C54), GM24143 (RRID:CVCL_1C48), GM24631 (RRID:CVCL_1C97), GM24694 (RRID:CVCL_1C98), and GM24695 (RRID:CVCL_1C99) at the Coriell Institute for Medical Research National Institute for General Medical Sciences cell line repository. The Genome in a Bottle Consortium selected these seven genomes for characterization because the pilot HG001 had extensive pre-existing public data, and HG002 to HG007 are two trios from the Personal Genome Project that have a broader consent that permits commercial redistribution and recontacting participants for further sample collection.

## Notes

https://ftp-trace.ncbi.nlm.nih.gov/ReferenceSamples/giab/release/

### Summary of Updates

fix author list

