## Supplementary Figs1-7, Tables 1,3-4,7,9-10,13-14, and Notes1-2 for "Benchmarking challenging small variants with linked and long reads"

**Supplementary Table 1**: Coverage of each sample by linked-reads and long-reads. PacBio HiFi calculated from median depth in VCFs used in integration pipeline. 10x Genomics coverage estimates are from the sequencing provider. ONT coverage calculated from median of mosdepth 1000 bp windows in bam file (note that variants from ONT were not used in v4.2.1; it was only used to exclude regions with abnormal coverage).

| | Reference | PacBio HiFi coverage | 10x coverage | ONT coverage |
|---|---|---|---|---|
| HG001 | GRCh37 | 68 | 75 | 37 |
| HG001 | GRCh38 | 67 | 75 | 37 |
| HG002 | GRCh37 | 54 | 84 | 59 |
| HG002 | GRCh38 | 54 | 84 | 59 |
| HG003 | GRCh37 | 62 | 71 | 84 |
| HG003 | GRCh38 | 63 | 71 | 85 |
| HG004 | GRCh37 | 60 | 69 | 85 |
| HG004 | GRCh38 | 60 | 69 | 85 |
| HG005 | GRCh37 | 47 | 53 | 57 |
| HG005 | GRCh38 | 47 | 53 | 59 |
| HG006 | GRCh37 | 67 | 53 | 51 |
| HG006 | GRCh38 | 67 | 53 | 51 |
| HG007 | GRCh37 | 56 | 53 | 41 |
| HG007 | GRCh38 | 56 | 53 | 41 |

**Supplementary Table 2:** Comparison of v4.2.1 to v3.3.2 using hap.py with v2.0 genome stratifications are available in SupplementaryTable2_HG002_GRCh37_v4.2.1_vs_v3.3.2.extended.xlsx

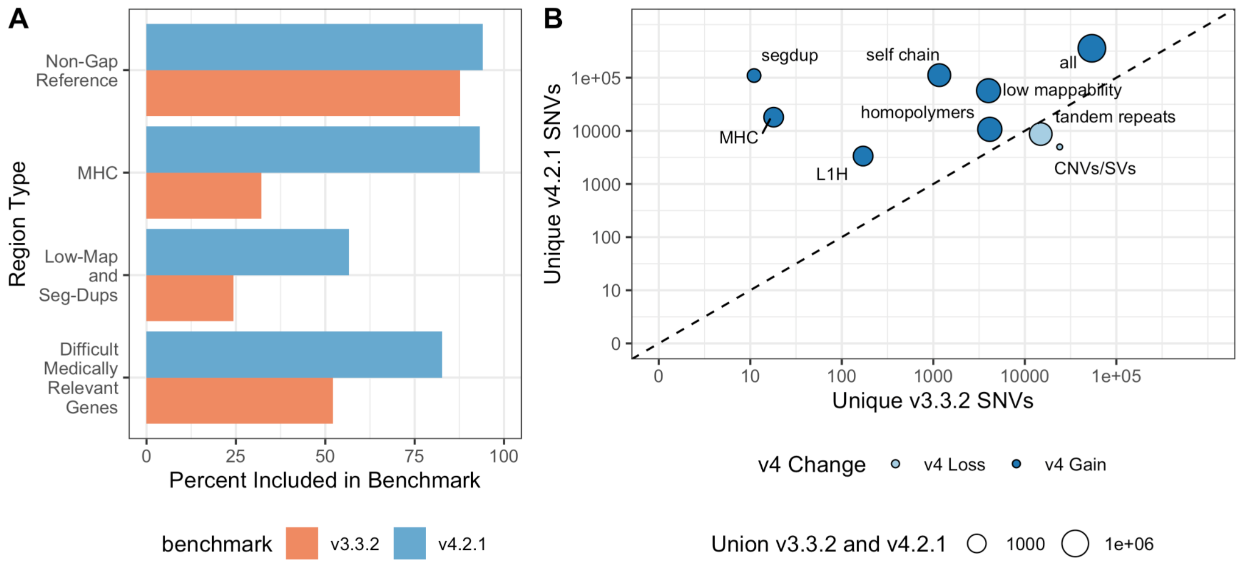


**Supplementary Figure 1**: New benchmark set for GRCh37 includes more of the reference genome and more SNVs and indels**.**

**Supplementary Table 3**: Errors in v3.3.2 identified in (Wenger et al. 2019) that are updated now matching PacBio HiFi callset or removed from benchmark regions.

| Chromosome | Position | Result | Region Type |
|---|---|---|---|
| 4 | 11,468,804 | Outside v4.2.1 benchmark regions | |
| 5 | 42,740,225 | Outside v.42.1 benchmark regions | LINE:L1PA2 |
| 2 | 5,143,996 | Call matches in benchmark region | |
| 13 | 48,291,499 | Outside v4.2.1 benchmark regions | LINE:L1PA3 |
| 8 | 5,930,728 | Outside v4.2.1 benchmark regions | |
| 15 | 41,943,823 | Outside v4.2.1 benchmark regions | |
| 6 | 9,737,425 | Outside v4.2.1 benchmark regions | |
| 7 | 157,385,671 | Reference call in benchmark regions | |
| 17 | 32,064,214 | Outside v4.2.1 benchmark regions | |
| 1 | 94,256,825 | Call matches in benchmark region | LINE:L1PA2 |
| 2 | 153,864,971 | Call matches in benchmark region | LINE:L1HS |
| 4 | 112,819,087 | Call matches in benchmark region | LINE:L1HS |
| 4 | 165,026,074 | Call matches in benchmark region | LINE:L1PA2 |

| | | | |
|---|---|---|---|
| 11 | 23,338,682 | Call matches in benchmark region | LINE:L1P1 |
| 1 | 35,034,071 | Call matches in benchmark region | LINE:L1HS |
| 3 | 79,181,734 | Call matches in benchmark region | LINE:L1HS |
| 4 | 94,532,444 | Call matches in benchmark region | LINE:L1HS |
| 8 | 46,873,565 | Outside v4.2.1 benchmark regions | |
| 9 | 22,350,168 | Call matches in benchmark region | LINE:L1PA2 |
| 21 | 42,288,851 | Call matches in benchmark region | LINE:L1PA2 |

**Supplementary Table 4:** Benchmark set overlap of 163 difficult-to-map, medically-relevant genes in GRCh37. There are 10,152,047 bps in GRCh37 for medically-relevant genes that are difficult to sequence for short reads genes on the primary assembly for chromosomes 1-22.

| Benchmark Set | bp included | SNVs | INDELS |
|---|---|---|---|
| v3.3.2 | 5,283,743 (52.0%) | 6,364 | 997 |
| v4.2.1 | 8,428,864 (83.0%) | 10,710 | 1,471 |

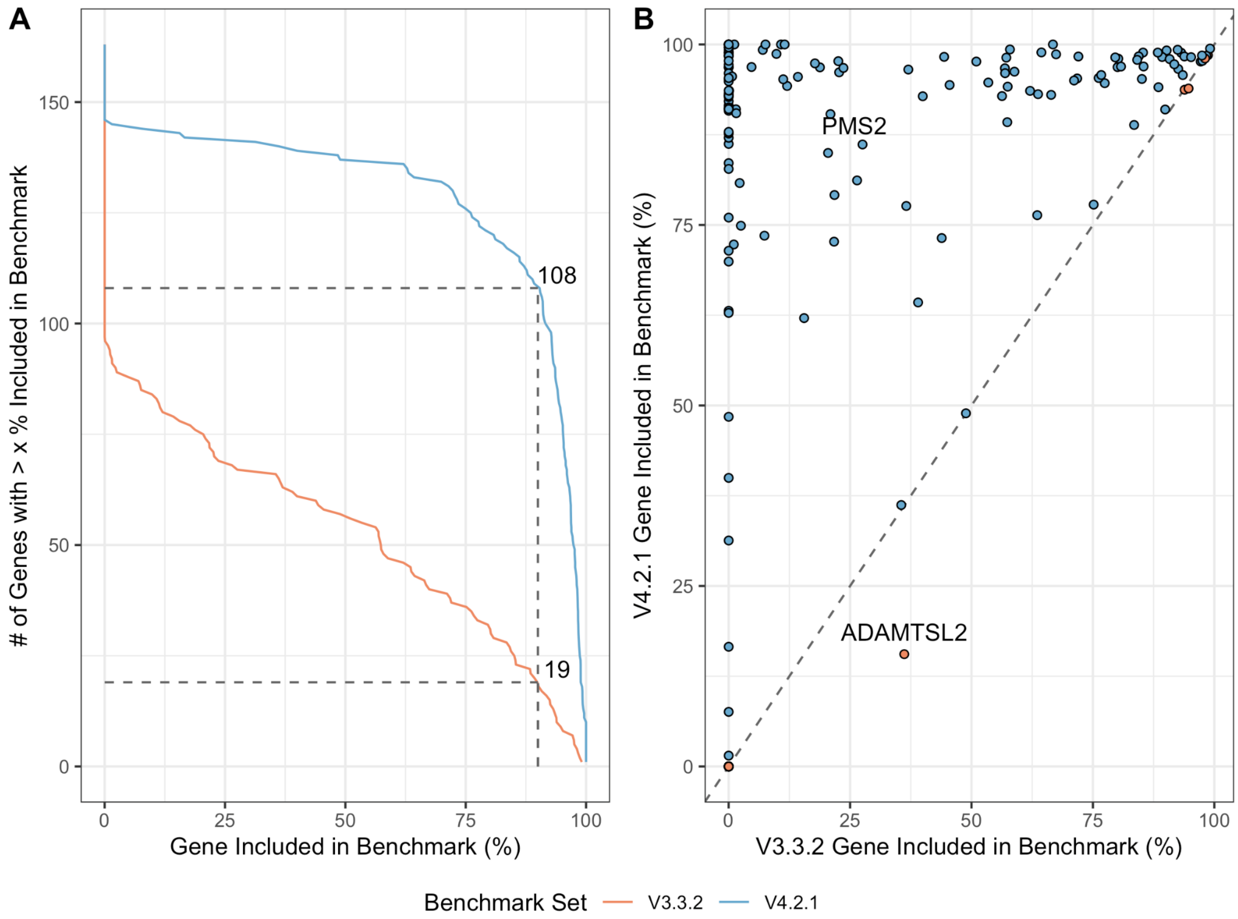


**Supplementary Figure 2: Included bases of difficult-to-map, medically-relevant genes by GRCh37 v4.2.1.**

**Supplementary Table 5: Detailed results of long range PCR and Sanger sequencing confirmation:** Available in SupplementaryTable5_LongRangePCRConfirmationResults.xlsx

**Supplementary Table 6: Included bases of each difficult-to-map, medically-relevant gene in v3.3.2 vs v4.2.1 for GRCh37 and GRCh38:** Available in SupplementaryTable6_difficult_medically_relevant_gene_cov.xlsx

**Supplementary Table 7**: Manual curation results of 10 random sites in HG002 v4.2.1 that match Category 1 SNVs in Platinum Genomes.

| Chromosome | Position | Curation | Notes |
|---|---|---|---|
| 16 | 18288432 | Selfchain/segdup | Many variants on one HP CCS, in selfchain/segdup |
| 19 | 54726776 | Selfchain/segdup | Cluster of variants in 10x/Illumina nearby and CCS has more variants on one HP than the other. In high depth selfchain/segdup that is smaller than 10kb |
| 19 | 41379908 | Selfchain/segdup | Cluster of variants, in LINE:L1MA3. In depth 2 segdup and depth 2 selfchain |
| 8 | 11872949 | Selfchain/segdup | Potential SV in segdup, since CCS and ONT have clipped reads nearby. Cluster of CAT1 variants |
| 15 | 20360478 | Possible CNV | Likely CNV given CCS data and high coverage in ONT. Several CAT1 variants in the region. |
| 8 | 7223157 | Selfchain/segdup | Cluster of CAT1 variants, in segdup and normal coverage but in a cluster of variants on one HP CCS and ONT, so may be more complex |
| 15 | 20453992 | Possible CNV | Cluster of CAT1 variants, cluster of variants on one HP CCS, and large change in coverage in CCS and nearby SV |
| 7 | 149749666 | Possible CNV | Cluster of CAT1 variants. Large changes in coverage in region in CCS data but overall looks reasonable. |
| 15 | 20454464 | Possible CNV | Many CAT1 variants in region, large change in CCS coverage in region, near what appears to be SV that is excluded from v4.2.1 |
| 12 | 74899879 | Possible CNV | Somewhat elevated CCS coverage. In LINE:L1PA3. |

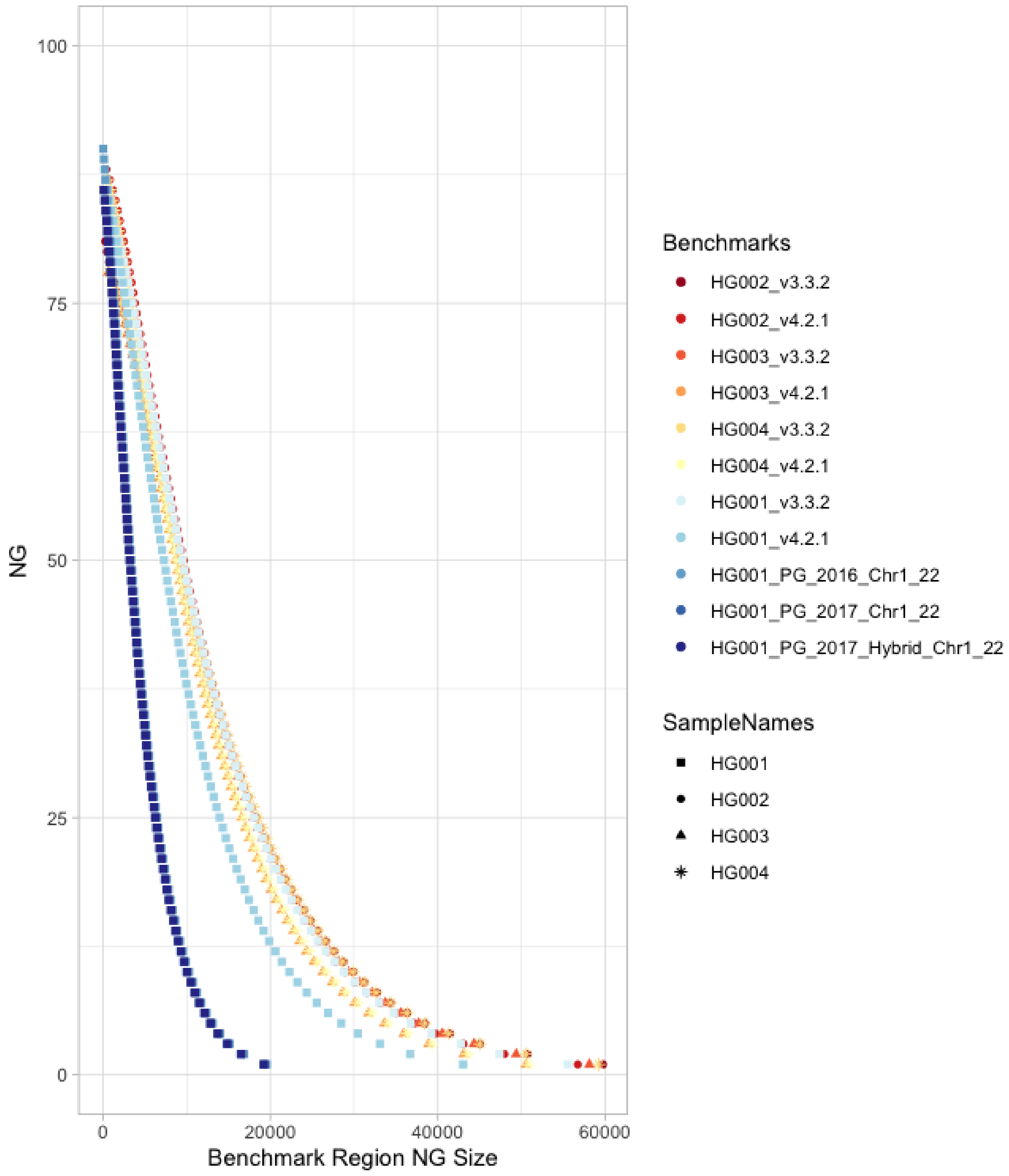


**Supplementary Figure 5**: NG is the percent of the GRCh38 reference covered by benchmark regions at least as large as the Benchmark Region NG Size. This metric is analogous to Assembly NG50 except that benchmark region size is used in place of contig length. The contiguity of the benchmark improves in v4.2.1 compared to v3.3.2 and all versions of Platinum Genomes (PG).

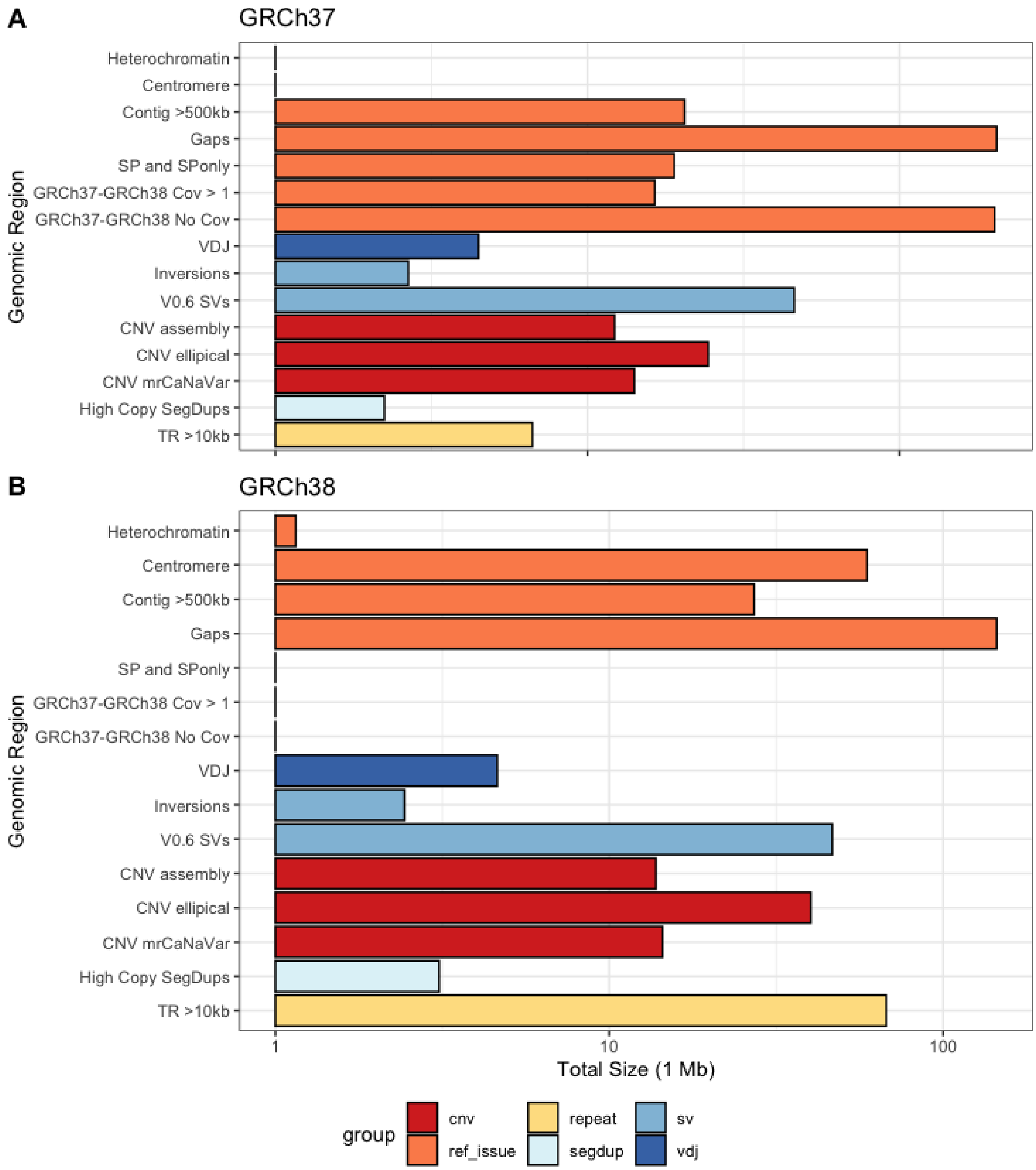


**Supplementary Figure 6**: **Base pairs in genomic regions excluded for all input variant call sets.**

**Supplementary Table 8:** Errors in v3.3.2 GRCh38 corrected hap.py results are available in SupplementaryTable8_GRCh38_v3.3.2_corrections.xlsx

**Supplementary Table 9:** Problematic regions in GRCh38 v3.3.2 that were near or in the centromere.

| GIAB Sample | Chromosome | Start | End | Region Type |
|---|---|---|---|---|
| HG002 | chr8 | 43637994 | 43672749 | Centromere |
| HG002 | chr8 | 43603010 | 43637285 | Centromere |
| HG002 | chr8 | 43831369 | 43864819 | Centromere |
| HG002 | chr7 | 62742402 | 62800702 | q11.21 (near centromere) |
| HG002 | chr7 | 57925899 | 57969199 | p11.1 (centromere) |
| HG002 | chr7 | 54317738 | 54350806 | p11.2 |
| HG002 | chr7 | 62821943 | 62851720 | q11.21 |
| HG002 | chr5 | 46337535 | 46371375 | Centromere |
| HG002 | chr5 | 46009909 | 46041150 | Centromere |
| HG002 | chr3 | 90613721 | 90676762 | Centromere |
| HG002 | chr3 | 90268364 | 90303792 | Centromere |
| HG002 | chr3 | 90445745 | 90478995 | Centromere |
| HG002 | chr19 | 27523978 | 27570562 | Centromere |
| HG002 | chr12 | 37624574 | 37664823 | Centromere |
| HG002 | chr12 | 34536432 | 34575253 | Centromere |
| HG002 | chr12 | 34483102 | 34520344 | Centromere |
| HG002 | chr12 | 37342974 | 37379851 | Centromere |
| HG002 | chr11 | 50785848 | 50821348 | p11.12 (near centromere) |
| HG002 | chr10 | 39052350 | 39083950 | Centromere |
| HG002 | chr10 | 39116363 | 39147923 | Centromere |
| HG003 | chr8 | 43601909 | 43637285 | Centromere |
| HG003 | chr8 | 43637994 | 43672749 | Centromere |
| HG003 | chr7 | 62742945 | 62800702 | q11.21 (near centromere) |
| HG003 | chr5 | 50193424 | 50229094 | Centromere |
| HG003 | chr5 | 46337535 | 46371375 | Centromere |
| HG003 | chr4 | 8843663 | 8892454 | p16.1 |
| HG003 | chr3 | 90598264 | 90676761 | Centromere |
| HG003 | chr3 | 90268364 | 90303792 | Centromere |
| HG003 | chr3 | 90411794 | 90445037 | Centromere |
| HG003 | chr3 | 90445745 | 90478939 | Centromere |
| HG003 | chr22 | 22145576 | 22178716 | q11.22 |

| | | | | |
|---|---|---|---|---|
| HG003 | chr19 | 27523978 | 27577850 | Centromere |
| HG003 | chr12 | 37624574 | 37664823 | Centromere |
| HG003 | chr12 | 34536383 | 34575253 | Centromere |
| HG003 | chr12 | 34482540 | 34520344 | Centromere |
| HG003 | chr12 | 37342974 | 37379851 | Centromere |
| HG003 | chr11 | 50772422 | 50821348 | p11.12 (near centromere) |
| HG003 | chr10 | 39013337 | 39083750 | Centromere |
| HG003 | chr10 | 39116363 | 39153579 | Centromere |
| HG003 | chr10 | 39183589 | 39216647 | Centromere |
| HG004 | chr8 | 43637994 | 43672749 | Centromere |
| HG004 | chr8 | 43831342 | 43864819 | Centromere |
| HG004 | chr7 | 62742945 | 62815024 | q11.21 |
| HG004 | chr7 | 57925899 | 57969199 | Centromere |
| HG004 | chr5 | 46009909 | 46041150 | Centromere |
| HG004 | chr5 | 50193424 | 50223736 | Centromere |
| HG004 | chr4 | 144161988 | 144192833 | q31.21 |
| HG004 | chr3 | 90445745 | 90478957 | Centromere |
| HG004 | chr3 | 90507640 | 90536550 | Centromere |
| HG004 | chr2 | 88861923 | 88891174 | p11.2 |
| HG004 | chr19 | 27523978 | 27559503 | Centromere |
| HG004 | chr12 | 37263197 | 37300537 | Centromere |
| HG004 | chr12 | 37342828 | 37379552 | Centromere |
| HG004 | chr12 | 63767401 | 63796912 | q14.2 |
| HG004 | chr12 | 37815851 | 37844396 | Centromere |
| HG004 | chr12 | 34492116 | 34520344 | Centromere |
| HG004 | chr11 | 50772422 | 50806467 | p11.12 |
| HG004 | chr10 | 39120199 | 39153579 | Centromere |
| HG004 | chr10 | 39055460 | 39087970 | Centromere |
| HG004 | chr10 | 39183589 | 39213934 | Centromere |

**Supplementary Table 10**: Sequencing technology, mapping or assembler method, and variant caller that was used to generate each evaluation call set. The names used in **Figure 6** are in the fourth column.

| **Sequencing Technology** | **Variant Caller** | **Mapper/Assembler** | **Figure 6 Name** |
|---|---|---|---|
| PacBio HiFi | DeepVariant | mm2 | PB DV-mm2 |
| PacBio HiFi | GATK4 | mm2 | PB GATK4-mm2 |
| PacBio HiFi | Clair | mm2 | PB Clair-mm2 |
| PacBio HiFi | DV | Duplomap | PB DV-Duplomap |
| PacBio HiFi | dipcall | WHDenovo | PB Dipcall-WHDenovo |
| Illumina PCR-Free TruSeq 2x250bp | Dragen | Dragen | Ill Dragen |
| Illumina PCR-Free TruSeq 2x250bp | Dragen | VG | Ill Dragen-VG |
| Illumina PCR-Free HiSeq 2x150bp | SevenBridges | SevenBridges Graph Aligner | Ill SevenBridges GRAF |
| Illumina PCR-Free HiSeq 2x150bp | xAtlas | NovoAlign | Ill xAtlas |
| Illumina PCR-Free NovaSeq 2x250bp | GATK | BWA | Ill GATK-BWA |
| 10x Genomics | LongRanger | LongRanger | 10x LongRanger |
| 10x Genomics | paftools | Aquila | 10x paftools-Aquila |
| ONT | Clair | mm2 | ONT Clair-mm2 |
| ONT | Clair | ngmlr | ONT Clair-ngmlr |

### Supplementary Note 1: Benchmark Evaluation Call set Generation

#### Variant callsets used in evaluation

##### *PacBio HiFi reads with GATK Haplotype Caller*

HG002 HiFi reads from three publicly available datasets were aligned to the GRCh37 and GRCh38 references using the pbmm2 v0.10.0 with `--preset CCS`. Small variants were called with GATK v4.0.10.1 HaplotypeCaller with `--pcr-indel-model AGGRESSIVE` and `--minimum-mapping-quality 10`. Variants were filtered on the QD (Quality by Depth) value with GATK v4.0.10.1 Variant Filtration, such that:

- SNVs with QD < 2 are filtered
- Indels > 1bp with QD < 2 are filtered
- 1 bp Indels with QD < 5 are filtered

| Instrument | Insert Size | SRA | FTP |
|---|---|---|---|
| Sequel I System | 10 kb | -- | ftp://ftp.ncbi.nlm.nih.gov/ReferenceSamples/giab/data/AshkenazimTrio/HG002_NA24385_son/PacBio_CCS_10kb |
| Sequel I System | 15 kb | SRX5327410 | ftp://ftp.ncbi.nlm.nih.gov/ReferenceSamples/giab/data/AshkenazimTrio/HG002_NA24385_son/PacBio_CCS_15kb |
| Sequel II System | 11 kb | SRX5527202 | ftp://ftp.ncbi.nlm.nih.gov/ReferenceSamples/giab/data/AshkenazimTrio/HG002_NA24385_son/PacBio_SequelII_CCS_11kb |

GRCh37 reference used for alignment:
ftp://ftp-trace.ncbi.nih.gov/1000genomes/ftp/technical/reference/phase2_reference_assembly_sequence/hs37d5.fa.gz
GRCh38 reference used for alignment:
ftp://ftp.ncbi.nlm.nih.gov/genomes/all/GCA/000/001/405/GCA_000001405.15_GRCh38/seqs_for_alignment_pipelines.ucsc_ids/GCA_000001405.15_GRCh38_no_alt_analysis_set.fna.gz
https://github.com/PacificBiosciences/pbmm2
https://github.com/broadinstitute/gatk/releases/tag/4.0.10.1

##### *PacBio Hifi reads using minimap2 with DeepVariant*

A set of ~80x coverage PacBio CCS data was mapped to each reference:

minimap2 VN:2.15-r905

minimap2 -ax asm20 -t 32

(Note that the mapping of these files predates some improved recommendations for mapping to use pbmm2)

DeepVariant v0.8 with the PACBIO model was applied to the mapped files. The commands and workflow used are identical to the DeepVariant case-study:

https://github.com/google/deepvariant/blob/r0.8/docs/deepvariant-pacbio-model-case-study.md

No filtering is applied.

##### *PacBio HiFi reads re-aligned using Duplomap*

HG002 HiFi reads aligned to the GRCh37 reference using Minimap2 were downloaded from ftp://ftp-trace.ncbi.nlm.nih.gov/ReferenceSamples/giab/data/AshkenazimTrio/HG002_NA24385_son/PacBio_CCS_15kb_20kb_chemistry2/ and reads overlapping segmental duplications were realigned using a tool Duplomap (https://gitlab.com/tprodanov/duplomap) that used paralogous sequence variants to map reads with multiple possible alignment locations.  Small variants were called from the realigned bam file using DeepVariant v.0.8 with default parameters.

##### *10x Genomics using Aquila local assembly*

Aquila uses linked-read data for generating a high quality diploid genome assembly, from which it then comprehensively detects and phases personal genetic variation. Here, Aquila merged two link-reads libraries to generate WGS variant calls for NA24385. Assemblies and VCFs for this merged library L5 + L6 can be found at http://mendel.stanford.edu/supplementarydata/zhou_aquila_2019/. The raw linked-reads fastq files can be downloaded in the Sequence Read Archive and its BioProject accession number is PRJNA527321.

##### *Illumina TruSeq DNA PCR-Free reads with Illumina Dragen Bio-IT platform*

Illumina PCR-Free reads (2 x 250bp with 350bp insert size) are downloaded from the public file server. Dragen 3.3.7 is used to perform alignment, variant calling, and filtering on GRCh37 and GRCh38 reference assemblies. Variant filtering is based on MQ (Mapping Quality), MQRankSum (Z-score From Wilcoxon rank sum test of Alt vs Ref read MQs), and ReadPosRankSum (Z-score from Wilcoxon rank sum test of Alt vs Ref read position bias) values. For SNVs, MQ < 30.0, MQRankSum < -12.5, or ReadPosRankSum < -8.0 are filtered out. For INDEL, ReadPosRankSum < -20.0 are filtered.

Illumina PCR-Free reads are downloaded from ftp://ftp-trace.ncbi.nlm.nih.gov/ReferenceSamples/giab/data/AshkenazimTrio/HG002_NA24385_son/NIST_Illumina_2x250bps/reads/

##### *Illumina TruSeq DNA PCR-Free reads with VG alignment and Illumina Dragen Bio-IT platform*

Illumina PCR-Free read pairs (2 x 250bp with 350bp insert size) are downloaded from and extracted from novoaligned bams that are hosted on the public file server. The process is based on aligning the HG002 to genome graphs that were constructed from HG003 and HG004 parental variants. All alignments are performed using Variation Graph Toolkit (VG) and variant calling is done using Dragen version 3.2. Default variant calling settings in Dragen 3.2 were used during GVCF and VCF variant calling. The methods used to convert graph alignments to linear alignments and parental graph construction are in the workflow defined on the vg_wdl GitHub repository.

The workflow used to process this data can be found here
https://github.com/vgteam/vg_wdl/blob/master/workflows/vg_trio_multi_map_call.wdl

Illumina PCR-Free reads for the trio used in parental graph construction and HG002 alignment are downloaded from
ftp://ftp-trace.ncbi.nlm.nih.gov/ReferenceSamples/giab/data/AshkenazimTrio/HG002_NA24385_son/NIST_Illumina_2x250bps/novoalign_bams/
ftp://ftp-trace.ncbi.nlm.nih.gov/ReferenceSamples/giab/data/AshkenazimTrio/HG003_NA24149_father/NIST_Illumina_2x250bps/novoalign_bams/
ftp://ftp-trace.ncbi.nlm.nih.gov/ReferenceSamples/giab/data/AshkenazimTrio/HG004_NA24143_mother/NIST_Illumina_2x250bps/novoalign_bams/

The population data used for initial graph alignments of the HG002 trio samples are based on the 1000 genomes phase 3 variant dataset and the GRCh37 reference genome.
http://ftp.1000genomes.ebi.ac.uk/vol1/ftp/release/20130502/ALL.wgs.phase3_shapeit2_mvncall_integrated_v5b.20130502.sites.vcf.gz

##### *10x genomics using LongRanger with GATK Haplotype Caller*

These callsets, generated independently for each individual in the Ashkenazi trio, used LongRanger[21] (version 2.2, code at https://github.com/10XGenomics/longranger) and GATK v4.0.0.0 as variant caller with default parameters on 10x Genomics linked-reads data for the family trio (84x, 70x, and 69x coverage for HG002 NA24385 son, HG003 NA24149 father, and HG004 NA24143 mother, respectively) against both GRCh37 and GRCh38. The vcf and bam files for each genome are under:
ftp://ftp-trace.ncbi.nlm.nih.gov/ReferenceSamples/giab/data/AshkenazimTrio/analysis/10XGenomics_ChromiumGenome_LongRanger2.2_Supernova2.0.1_04122018/

The variant curation used the 10x Genomics VCF from LongRanger 2.2 (SRA accession SRX2225480), which is available at:
ftp://ftp-trace.ncbi.nlm.nih.gov/ReferenceSamples/giab/data/AshkenazimTrio/analysis/10XGenomics_ChromiumGenome_LongRanger2.2_Supernova2.0.1_04122018/GRCh37/NA24385_300G/NA24385.GRCh37.phased_variants.vcf.gz

All samples were sequenced on the Illumina Xten at 2x150bp. The Ashkenazim trio was done using the v1 of the 10x library prep protocol.

##### *HiFi Clair*

This callset was generated using Sequel II 11kbp HiFi reads aligned to the hs37d5 reference with pbmm2, publicly available here: ftp://ftp-trace.ncbi.nlm.nih.gov/ReferenceSamples/giab/data/AshkenazimTrio/HG002_NA24385_son/PacBio_SequelII_CCS_11kb/. The variants were called by using Clair (v1) on these alignments.

##### *Illumina Novaseq 2x250bp data*

*The sample HG002 was sequenced on an Illumina Novaseq 6000 instrument with 2x250bp paired end reads at the New York Genome Center. The libraries were prepped using Truseq DNA PCR-free library preparation kit. The raw reads were aligned to both GRCh37 and GRCh38 human reference. Alignment to GRCh38 reference, marking duplicates and base quality recalibration was performed as outlined in the Centers for Common Disease Genomics (CCDG) functional equivalence paper. Alignment to GRCh37 was performed using BWA-Mem (ver. 0.7.8) and marking duplicates using Picard (ver. 1.83) and local Indel realignment and base quality recalibration using GATK (ver. 3.4-0). Variant calling was performed using GATK (ver. 3.5) adhering to the best practices recommendations from the GATK team. Variant calling constituted generating gVCF using HaplotypeCaller, genotyping using the GenotypeGVCFs subcommand and variant filtering performed using VariantRecalibrator and ApplyRecalibration steps. A tranche cutoff of 99.8 was applied to SNP calls and 99.0 to InDels to determine PASS variants.*

*The raw reads are available for download at SRA.*
https://www.ncbi.nlm.nih.gov/sra/SRX7925517[accn]
https://www.ncbi.nlm.nih.gov/sra/SRX7925518[accn]
https://www.ncbi.nlm.nih.gov/sra/SRX7925519[accn]

**Supplementary Table 11:** Manual Curation Results are available in SupplementaryTable11_v4.1_ManualCurationResults.xlsx

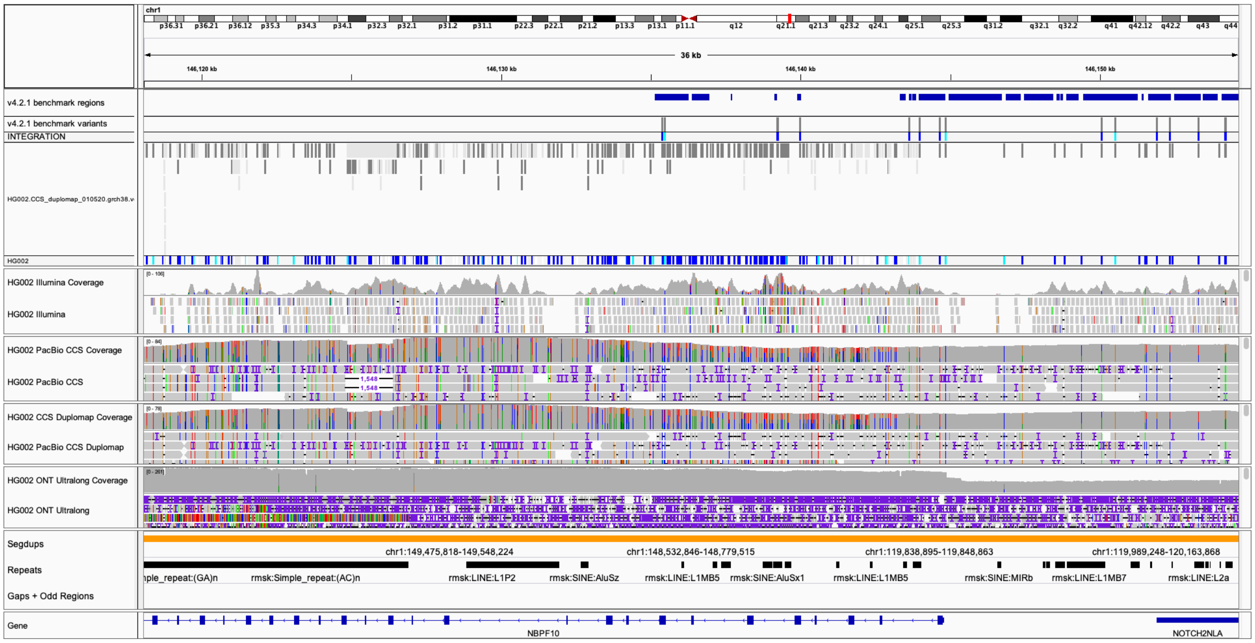


**Supplementary Figure 7:** The benchmark and callset both make calls in this region where there is likely a large duplication in HG002 compared to GRCh38 that was not detected by our exclusion criteria. This specific example is a FP SNP in PacBio HiFi Duplomap DV where the benchmark region indicates a reference call at this location.

### Supplementary Note 2: Process from v4alpha to v4.2.1

The Genome in a Bottle Consortium has an iterative evaluation process to ensure new benchmarks are useful for assessing performance across diverse sequencing technologies and variant calling methods. The first version using PacBio HiFi and 10x Genomics data was v4alpha. In particular, GIAB found that some regions contained unreliable calls across technologies, and these regions were excluded from subsequent releases. In addition, the PacBio HiFi data changed during releases as new data were collected, and the 10x Genomics data remained constant at ~84x coverage.

The v4alpha release used PacBio Sequel I HiFi ~15 kb reads at ~28x coverage, with the difficult regions below (Bold entries changed from v3.3.2):

| Difficult Region Description | Method Excluded From |
|---|---|
| All candidate structural variant regions from the Son-Mother-Father Trio | All methods |
| All tandem repeats < 51bp in length | All methods except GATK from Illumina PCR-free, Complete Genomics, and **PacBio CCS DeepVariant** |
| All tandem repeats > 51bp and < 200bp in length | All methods except GATK from Illumina PCR-free and **PacBio CCS DeepVariant** |
| All tandem repeats > 200bp in length | All methods except **PacBio CCS DeepVariant** |
| Perfect or imperfect homopolymers > 10bp | All methods except GATK from Illumina PCR-free |

| | |
|---|---|
| Segmental duplications from Eichler *et al*. | All methods except **10X Genomics and PacBio CCS** |
| Segmental duplications > 10Kbp from self-chain mapping | All methods except **10X Genomics and PacBio CCS** |
| Regions homologous to contigs in hs37d5 decoy | All methods except **10X Genomics and PacBio CCS** |
| Difficult to map regions for short reads | All methods except **10X Genomics and PacBio CCS** |
| Homopolymer > 6bp in length | **All methods except GATK from Illumina PCR-free and Complete Genomics** |

The v4beta release used PacBio Sequel I HiFi ~15 kb reads at ~28x coverage. Additionally, v4beta used additional tandem repeat files from UCSC, excluded the entire tandem repeat if any part was not in the benchmark BED, and changed the difficult regions below:

| Difficult Region Description | Method Excluded From |
|---|---|
| v0.6 SV Benchmark | All methods |
| Regions that are collapsed and expanded from GRCh37/38 Primary Assembly Alignments (corrected) | All methods |
| Diploid assemblies exhibit more than 2 contigs aligned > 10kb | All methods |
| Intersected short and long read based copy number > 2.5 (updated) | All Methods |
| Segmental duplications > 10Kb, Identity > 99%, Count > 5 | All methods |
| mrCaNaVar duplications > 10kb (052119) | All methods except 10X Genomics and PacBio CCS |
| Outliers from long read coverage | All Methods |
| LINE:L1Hs > 500 | All methods except Illumina MatePair, 10X Genomics, and PacBio CCS |
| All Tandem Repeats > 10kb in length | All methods |

The v4.0 release used PacBio Sequel II HiFi ~11 kb reads at ~32x coverage, updated to DeepVariant v0.8.

The v4.1 release used PacBio Sequel II HiFi ~15 kb and ~20 kb reads at ~52x coverage. The diploid assembly-based MHC benchmark was used for the MHC region in v4.1. We also added the difficult regions below:

| Difficult Region Description | Method Excluded From |
|---|---|
| **Potential copy number variation including CCS and ONT outlier and CCS, ONT, mrCanavar intersection** | **All methods** |
| **VDJ** | **All methods** |
| **Inversions** | **All methods** |

The v4.2 release is the first for HG003 and HG004, and it used hifiasm to perform the assembly of PacBio HiFi reads in the MHC, and used dipcall with this assembly to call variants, including in segmental duplications that were previously not assembled properly. Since it represents complex variants as individual SNVs and indels, dipcall helps improve partial credit in some cases for variants that are only partially called correctly by the query callset. We also excluded entire homopolymers and tandem repeats in the MHC if they were not completely covered by the benchmark bed. Since calls were made for HG003 and HG004 in addition to HG002, we also performed a trio Mendelian analysis and excluded Mendelian violations from the benchmark regions for all individuals (except putative de novo variants in HG002 were not excluded from the benchmark regions).

The v4.2.1 release is for HG002, HG003, and HG004 on both GRCh37 and GRCh38. We now use the same MHC hifiasm approach for HG002 as with HG003 and HG004. We exclude SVs from a pbsv call set from HG003 and HG004 in addition to the GIAB v0.6 SV benchmark. A final update is that we exclude the KIR region because of highly variable copy number.

**Supplementary Table 12:** Primer Sequences for Long-Range PCR are available in SupplementaryTable12_PrimerSequences.xlsx

**Supplementary Table 13: Long Range PCR Components**

| | ***LINEs*** | ***C4A*** | ***C4B*** | ***Cyp21A2*** | ***Cyp2D6*** | ***DMBT1*** | ***HSPG2*** | ***PMS2*** | ***STRC*** | ***TnxA*** | ***TnxB*** |
|---|---|---|---|---|---|---|---|---|---|---|---|
| Buffer (5X) | 1X | 1X | 1X | 1X | 1X | 1X | 1X | 1X | 1X | 1X | 1X |
| dNTP (250uM each) | 250uM | 400uM | 400uM | 250uM | 0.3mM | 400uM | 200uM | 400uM | 400uM | 250uM | 250uM |
| Forward Primer | 0.25uM | 0.5uM | 0.5uM | 10uM | 0.5uM | 0.4uM | 0.3uM | 0.2uM | 0.4uM | 10uM | 10uM |
| Reverse Primer | 0.25uM | 0.5uM | 0.5uM | 10uM | 0.5uM | 0.4uM | 0.3uM | 0.2uM | 0.4uM | 10uM | 10uM |
| Polymerase (1.25 units/uL) | 1.25 U | 1.25 U | 1.25 U | 1.25 U | 1.25 U | 2.5 U | 0.5 U | 1.25 U | 2 U | 0.5 U | 0.5 U |
| DNA | 300ng | 100ng | 100ng | 250ng | 1uL | 2uL | 300ng | 100ng | 300ng | 250ng | 250ng |
| Water | To 50uL | To 50uL | To 50uL | To 30uL | To 25uL | To 50uL | To 50uL | To 25uL | To 50uL | To 30uL | To 30uL |

**Supplementary Table 14**: **Long Range PCR Conditions**

| Gene | PCR Conditions |
|---|---|

| | |
|---|---|
| LINEs | 30 cycles of 98℃ for 10 seconds, 60℃ for 15 seconds, and 68℃ for 8 minutes. |
| *C4A* | 98℃ for 2 minutes; followed by 40 cycles of 98℃ for 45 seconds, 66℃ for 60 seconds, and 72℃ for 9 minutes, with a final extension step of 72℃ for 10 minutes. |
| *C4B* | 98℃ for 2 minutes; followed by 8 cycles of 94℃ for 45 seconds, 64℃ for 60 seconds, with a decrease of 0.5℃ per cycle, and 72℃ for 9 minutes; followed by 30 cycles of 94℃ for 45 seconds, 59℃ for 60 seconds, and 72℃ for 9 minutes, with an increase of 10 seconds per cycle, with a final extension step of 72℃ for 15 minutes |
| *Cyp21A2*<br>*TnxA*<br>*TnxB* | 94℃ for 4 minutes; followed by 12 cycles of 94℃ for 30 seconds, 62℃ for 40 seconds, and 68℃ for 5 minutes; followed by 16 cycles of 94℃ for 30 seconds, and 68℃ for 5 minutes. |
| *Cyp2D6* | 96℃ for 30 seconds; followed by 30 cycles of 94℃ for 15 seconds, 68℃ for 30 seconds, and 68℃ for 7 minutes, with a final extension step of 68℃ for 30 minutes. |
| *DMBT1* | 94℃ for 1 minute; followed by 30 cycles of 98℃ for 10 seconds, and 68℃ for 15 minutes, with a final extension step of 72℃ for 10 minutes. |
| *HSPG2* | 30 cycles of 98℃ for 10 seconds, 60℃ for 15 seconds, and 68℃ for 10 minutes. |
| *PMS2* | 94℃ for 1 minute; followed by 35 cycles of 94℃ for 10 seconds, and 65℃ for 30 seconds, and 68℃ for 15 minutes, with a final extension step of 72℃ for 10 minutes. |
| *STRC* | 93℃ for 3 minutes; followed by 38 cycles of 93℃ for 15 seconds, 64℃ for 30 seconds, and 68℃ for 17 minutes, with a final extension step of 68℃ for 5 minutes. |
